# Early development and co-evolution of microstructural and functional brain connectomes: A multi-modal MRI study in preterm and full-term infants

**DOI:** 10.1101/2024.09.09.612067

**Authors:** Gondová Andrea, Neumane Sara, Arichi Tomoki, Dubois Jessica

## Abstract

**Introduction:** Functional networks characterised by coherent neural activity across distributed brain regions have been observed to emerge early in neurodevelopment. Synchronized maturation across regions that relate to functional connectivity (FC) could be partially reflected in the developmental changes in underlying microstructure. Nevertheless, covariation of regional microstructural properties, termed ‘microstructural connectivity’ (MC), and its relationship to the emergence of functional specialization during the early neurodevelopmental period remains poorly understood.

**Methods:** We investigated the evolution of MC and FC postnatally across a set of cortical and subcortical regions, focusing on 45 preterm infants scanned longitudinally, and compared to 45 matched full-term neonates as part of the developing Human Connectome Project (dHCP) using direct comparisons of grey-matter connectivity strengths as well as network-based analyses.

**Results:** Our findings revealed a global strengthening of both MC and FC with age, with connection-specific variability influenced by the connection maturational stage. Prematurity at term-equivalent age was associated to significant connectivity disruptions, particularly in FC. During the preterm period, direct comparisons of MC and FC strength showed positive linear relationship, which seemed to weaken with development. On the other hand, overlaps between MC-and FC-derived networks (estimated with Mutual Information) increased with age, suggesting a potential convergence towards a shared underlying network structure that may support the co-evolution of microstructural and functional systems.

**Conclusion:** Our study offers novel insights into the dynamic interplay between microstructural and functional brain development and highlights the potential of MC as a complementary descriptor for characterizing the brain network development and alterations due to perinatal insults such as premature birth.

**Keypoints:**

1. Our study reveals a significant positive linear relationship between grey-matter functional connectivity and underlying microstructural connectivity during development, that decreases with age and varies across connection types.
2. Despite progressive maturational decoupling of microstructural and functional connectivity, a shared network structure may underlie changes in both properties.
3. Prematurity impacts the maturation of connectivity in both modalities, but with a higher reduction of functional than microstructural connectivity strengths.

## Introduction

Brain development during the third trimester of pregnancy and the perinatal period is characterized by a series of complex inter-related mechanisms. The resulting macro-and microstructural changes are crucial for establishing the structural and functional brain networks that support neurodevelopment and optimal outcomes (Gilmore et al., 2018). Recent advances in magnetic resonance imaging (MRI) have provided unprecedented access to study the brain during this critical period *in vivo* (Dubois et al., 2021).

In particular, exploring functional connectivity (FC) across cortical and subcortical brain regions with resting-state functional MRI (rs-fMRI) has revealed a strengthening of cortico-subcortical and cortico-cortical connectivity with distinct developmental patterns across various functional networks (Doria et al., 2010; Fransson et al., 2009; Jakab et al., 2014; Smyser et al., 2010; Taymourtash et al., 2023; van den Heuvel et al., 2015; Williams et al., 2023). By term-equivalent age (TEA), the topological architecture of FC partly resembles that of adults, with connectivity hubs observed in the earlier developing primary sensory and motor regions (Dall’Orso et al., 2022; Eyre et al., 2021; Fransson et al., 2009; Toulmin et al., 2015; Turk et al., 2019; van den Heuvel & Hulshoff Pol, 2010). Postnatally, FC maturation seems to progress asynchronously, following a primary-to-higher function order from sensorimotor/auditory to associative and default-mode networks (Cao et al., 2017; Eyre et al., 2021; Gao et al., 2015; Hoff et al., 2013).

This maturational progression seems consistent with the sequence of spatiotemporal maturation of grey matter (GM) microstructure taking place earliest in primary sensory regions, then association areas and prefrontal cortices as described by measures derived from quantitative structural MRI (Ball et al., 2013; Lebenberg et al., 2019; Monson et al., 2018; Neil & Smyser, 2018; Yu et al., 2016). Thus, investigating the covariation of microstructural descriptors across the GM regions of interest (ROIs), i.e. *microstructural connectivity* (MC), and its relationship to emerging patterns of FC could reveal synchrony of microstructural features across cortical and subcortical regions belonging to the same developing functional network (Alexander-Bloch et al., 2013), and provide insights into coordinated maturation across different brain modalities. While previous studies have explored functional and microstructural developmental changes within emerging brain networks separately, this study expands on the previous work to directly investigate their relationship during the preterm period across various cortical GM ROIs. We hypothesize that early microstructural connectivity serves as a foundation for the development of functional connectivity, with distinct maturation profiles observable across different subsets of connections depending on their maturational trajectories.

The idea that the covariation of GM features can be interpreted as biologically meaningful and functionally relevant units was first proposed in foundational early studies based on histological assessments of cortical cytoarchitecture (Brodmann, 1908; von Economo & Koskinas, 1925). More recent anatomical MRI studies that model the covariation of morphometric markers across cortical regions (such as cortical thickness that indirectly reflects the underlying microstructure) have further supported this notion (Alexander-Bloch et al., 2013; King & Wood, 2020), highlighting a higher likelihood of anatomical connectivity between morphologically similar brain regions (Barbas, 2015; Goulas et al., 2016, 2017; Seidlitz et al., 2018), as well as similarities in their genetic and transcriptomic profiles (Alexander-Bloch et al., 2013; Yee et al., 2018).

Importantly, these regional covariations are sensitive to neurodevelopmental and age-related changes (Khundrakpam et al., 2013, 2016; Romero-Garcia et al., 2018; Váša et al., 2018), with groups of regions showing similar morphometric profiles and developmental trajectories (Alexander-Bloch et al., 2013). Despite potential interpretations, only a few studies focused on the first 2 postnatal years to explore developmental relationships of regional covariation based on markers such as GM volume (Fan et al., 2011), cortical thickness (Geng et al., 2016; Nie et al., 2014), cortical folding (Nie et al., 2014), and fibre density (Fan et al., 2011; Nie et al., 2014). However, these studies led to highly heterogenous results, likely due to the employment of single descriptors with specific spatial and temporal developmental patterns that might influence the estimated relationships (Gilmore et al., 2012; Lyall et al., 2015; Nie et al., 2014; Seidlitz et al., 2018). Recent multiparametric approaches have integrated multiple morphological and microstructural descriptors derived from diffusion MRI (dMRI) using diffusion models such as Diffusion Tensor Imaging (DTI) (Basser et al., 1994) and Neurite Orientation Dispersion and Density Imaging (NODDI) (Zhang et al., 2012). These models provide complementary measures sensitive to changes in neuronal and glial density, neurite complexity, synaptic overproduction and pruning, and reduction in brain water content (Ouyang, Dubois, et al., 2019) that can be combined to provide more comprehensive description of the underlying microstructural developmental changes within the GM. Resulting estimates of structural covariance in the neonatal brain were used to delineate modules consistent with known cytoarchitectonic tissue classes and functional systems (Fenchel et al., 2020), and to allow prediction of social-emotional performance at 18 months in full-term (FT) newborns (Fenchel et al., 2022) and the discrimination of preterm (PT) and FT individuals at TEA (Galdi et al., 2020).

Expanding on these works, here we investigate the relationships between microstructural and functional development in the preterm period using multimodal (anatomical, diffusion, and resting-state functional) MRI data from the developing Human fitConnectome Project (dHCP) (Edwards et al., 2022) to analyse 45 preterm-born infants scanned twice (near birth: PT:ses1, median postmenstrual age (PMA) at scan 34.9 weeks, range [28.3w–36.9w]; and close to term-equivalent age (TEA): PT:ses2, median PMA at scan 41.3 weeks, range [38.4w–44.9w]), and 45 full-term control neonates matched for PMA at scan (with PT:ses2) and sex. At the methodological level, in contrast to previous subject-level multiparametric studies (Fenchel et al., 2020; Galdi et al., 2020), we employed a group-wise approach to account for the reduced number of metrics we employed for microstructural similarity estimation and the need of corrections for confounders such as gestational age (GA) at birth required for the group comparisons.

Changes in FC and MC between the infant groups were evaluated for each modality separately before describing the MC-FC relationship. Since disruptions of normal gestation, such as preterm birth, can lead to significant heterogenous and region-specific alterations in both GM microstructural maturation (Ball et al., 2013; Batalle et al., 2019; Dimitrova et al., 2021; Eaton-Rosen et al., 2015, 2017; Mukherjee et al., 2001; Ouyang, Jeon, et al., 2019; Smyser et al., 2016; Yu et al., 2016) and functional connectivity (Ball et al., 2016; Brenner et al., 2021; Keunen et al., 2017; Smyser et al., 2010), we also assessed deviations related to prematurity (PT:ses2 vs FT) in both modalities.

To test the hypothesis that early developing microstructural clusters and connections might serve as the foundation for synchronised maturation across brain areas and thus efficient functionality of brain networks (Alexander-Bloch et al., 2013),we attempted to investigate the potential direction of the early MC-FC relationship in a longitudinal manner in the preterm group. Additionally, to assess whether some connections might show distinct MC-FC maturation profiles and dynamics in the perinatal period, our analyses focused on specific subsets of ROI connections with expected maturational differences as previously described for functional (van den Heuvel et al., 2015) and white matter structural (Kostović et al., 2019) connectivity: cortico-subcortical connections (including thalamo-cortical connections that were also highlighted separately given their crucial role in the preterm period (Kostović et al., 2021)); cortico-cortical connections (grouped as i) intra-hemispheric, inter-hemispheric ii) homotopic and iii) non-homotopic connections), anticipating potentially similar meaningful grouping at the level of MC. Connections involving the primary sensorimotor and visual ROIs were also highlighted given the early and intense development of these functions in the neonatal period. We further complemented our analyses with a network-based approach that involved hierarchical clustering of group MC and FC matrices. This allowed us to extend the direct MC-FC comparisons that could uncover progressive refinement and possible convergence of network structures between the two modalities, even in absence of direct relationships that could be related to different maturational stages of evaluated connections. The methodology of the present study is summarized in *Figure 1*.

**Figure 1.**
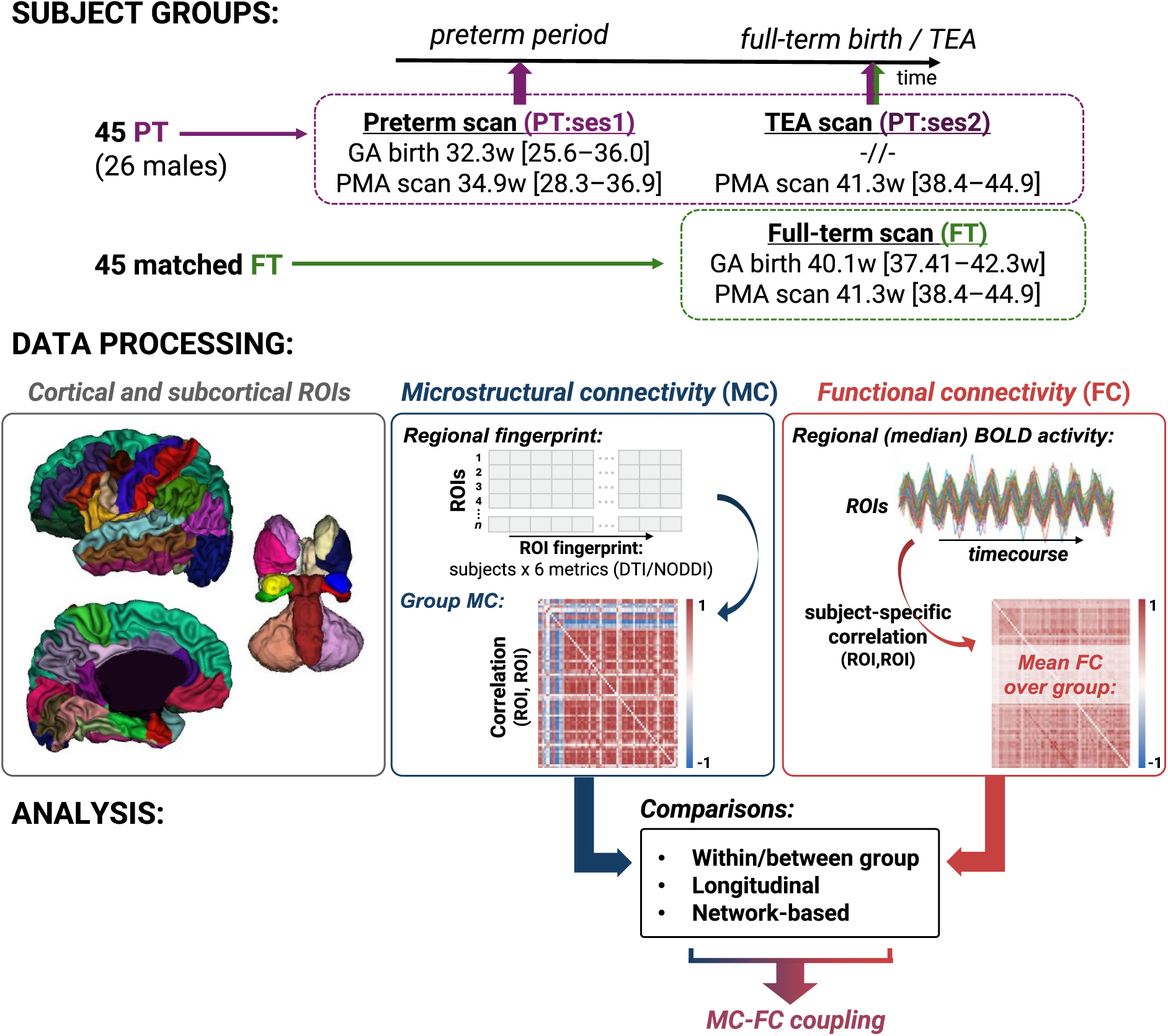
General analysis pipeline of the presented study.

## Materials and methods

### Data presentation

#### Subjects

This study included a sample of preterm and full-term neonates from the developing Human Connectome Project (dHCP) cohort (Edwards et al., 2022), collected at St Thomas’ Hospital London, UK from 2015 to 2020. This project received UK NHS research ethics committee approval (14/LO/1169, IRAS 138070), and written informed consent was obtained from the parents of all participant infants.

From the overall cohort, we identified 45 PT infants (26 males, median GA at birth 32.3 weeks, range [25.6w–36.0w]) who were scanned at two time points and whose dMRI and rs-fMRI data passed the quality control as described in 3^rd^ dHCP release notes. For the session 1, the infants were scanned in the preterm period at median postmenstrual age at scan of 34.9 weeks, range [28.3w–36.9w]; median birth-scan delay: 1.7 weeks, range [0.1w–9.3w]. For the session 2, infants were scanned close to TEA (median PMA at scan 41.3 weeks, range [38.4w–44.9w]; median birth-scan delay: 9.1 weeks, range [3.6w–15.6w]; median Ses1-Ses2 delay 7.3 weeks, range [2.7w–11.9w]). Note that PMA at Ses1 vs Ses2 were not correlated (Pearson’s r=0.18, p=0.17). Additionally, we considered a group of 45 FT infants matched to the preterm population on sex and age at MRI at TEA (GA at birth: median 40.1w, range [37.4w–42.3w]; median birth-scan delay: 0.4 weeks, range [0.1w–3.9w]). All included infants were without major brain focal lesions or any overt abnormality of clinical significance on anatomical MRI as evaluated by an expert paediatric neuroradiologist, (i.e., dHCP radiological scores were in the range [1-3]). More subject details for all three groups are available in *Supp. Figure 1.1*.

#### Acquisition and preprocessing of MRI data

MRI data were acquired using a Philips 3 Tesla Achieva scanner (Philips Medical Systems, Best, Netherlands). All infants were scanned during natural sleep using a neonatal head coil and imaging system optimized for the dHCP study as previously described (Hughes et al., 2017). In this study, we considered anatomical, diffusion, and resting-state functional MRI data available in its pre-processed state from the dHCP database (3^rd^ release) (Edwards et al., 2022).

The *anatomical data* resulted from acquisition and reconstruction using optimized protocols (Cordero-Grande et al., 2019), leading to super-resolved T2w images with an isotropic spatial voxel size of 0.5 mm. Processing followed a dedicated pipeline for segmentation and cortical surface extraction for T2w images of neonatal brains (Makropoulos et al., 2018), with bias-correction, brain extraction, volumetric segmentation using Draw-EM (Developing brain Region Annotation with Expectation Maximization) algorithm (Makropoulos et al., 2014), and reconstruction of white matter surface (inner cortical surface) meshes. These anatomical data were used for the extraction of GM ROIs (see section *Delineation of ROIs*).

Acquisition and reconstruction of the *diffusion data* (dMRI) followed a multi-shell high angular resolution diffusion imaging (HARDI) protocol with 4 b-shells (b = 0 s/mm2: 20 repeats; and b = 400, 1,000, 2,600 s/mm2: 64, 88, and 128 directions, respectively) (Hutter et al., 2018) and was pre-processed with correction for motion artifacts and slice-to-volume reconstruction using the SHARD approach, leading to an isotropic voxel size of 1.5 mm (Christiaens et al., 2021). Pre-processed data were used for the fitting of diffusion models and the measure of GM microstructure (see section *GM microstructural connectivity*).

*Resting state functional data* (rs-fMRI) was acquired for 15 minutes using a high temporal resolution multiband EPI protocol (TE=38 ms; TR=392 ms; MB factor=9x; 2.15 mm isotropic) (Price, 2015) and was processed following an automated processing framework specifically developed for neonates (Fitzgibbon et al., 2020). Available data was used for the estimation of the whole-brain functional connectivity (see section *Functional connectivity*).

More information on quality of the employed dMRI and rs-fMRI data can be found in the *Supp. Figure 2.1 & Supp. Table 2.1*.

### Estimation of connectivity matrices

#### Delineation of ROIs

Firstly, ROIs were defined as subregions of the cortical and subcortical grey matter to provide a framework for a focused and potentially interpretable assessment of the brain connectivity. Anatomically-driven parcellation strategy was used to provide a more comparable region correspondence between subjects and to allow the direct comparison of results between dMRI and rs-fMRI modalities. To define the ROIs, pre-processed anatomical data was used to parcellate the GM. Cortical parcels were defined on the cortical surface of each hemisphere using the M-CRIB-S surface-based parcellation tool optimized for the term-born neonates (Adamson et al., 2020) whose labelling scheme replicates the Desikan-Killiany-Tourville (DKT) atlas (Klein & Tourville, 2012). The subcortical ROIs were defined using a volumetric GM parcellation based on Draw-EM algorithm segmentation (Makropoulos et al., 2014), and included medial brainstem (bstem), and for each hemisphere: thalamus (thal, fusing high and low intensity regions), caudate nucleus (caud), lenticular nucleus (lenti), amygdala (amyg), hippocampus (hippo), and cerebellum (cereb). The 75 cortical and subcortical ROIs were combined and aligned to the subject diffusion and functional space with FSL 6.0’s FLIRT using precomputed warps provided within the dHCP database. The list and visualisation of ROIs used in this work is detailed in *Supp. Figure 2.2a*.

Because the M-CRIB-S approach was developed for full-term neonates, visual inspection of the ROI segmentation quality was performed on the 25 youngest PT infants at scan. While we observed an expected trend of an increase in the segmentation quality with PMA at scan (errors for the youngest subjects could be explained by the landmarks missing or being less pronounced, i.e., for example secondary and tertiary sulci), the parcellations remained satisfactory enough so as not to exclude any additional infants. Examples of the ROI longitudinal segmentations are shown in *Supp. Figure 2.2b*.

#### Microstructural connectivity (MC)

The DTI model was fitted to the diffusion data using a single shell (b = 1,000 s/mm2) and calculated with FSL’s DTIFIT to estimate metric maps for 4 metrics: fractional anisotropy (FA), axial diffusivity (AD), radial diffusivity (RD), and mean diffusivity (MD). Additionally, multi-shell diffusion data was used to derive the neurite density index (NDI) and orientation dispersion index (ODI) maps from the NODDI model (Zhang et al., 2012) using the CUDA 9.1 Diffusion Modelling Toolbox (cuDIMOT) NODDI Watson model implementation for GPUs (Hernandez-Fernandez et al., 2019). Derived NODDI maps were then corrected as described in (Neumane et al., 2022).

To create subject-specific regional microstructural fingerprints, we extracted median diffusion metrics for each cortical and subcortical ROI (*Supp. Materials: SI3. Univariate analyses of GM microstructure*). Beforehand, volumetric parcellations for subcortical ROIs underwent 1-voxel erosion to address potential border parcellation errors with surrounding white matter and cerebro-spinal fluid. For cortical ROIs, diffusion metrics were projected to the white-grey matter surface using a cylindrical approach guided by the minimum of AD, as described in (Lebenberg et al., 2019) and (Gondová et al., 2023). Four hemispheres with locally imperfect projections in the superior frontal gyrus were identified, but the error impact on median computations within such a large region was minimal.

To focus on the microstructural variability between ROIs, we aimed to correct for potential confounding factors influencing dMRI metrics and the resulting correlations between pairs of ROIs before computing the group connectivity matrices. Regional metric values were corrected independently over the 3 infant groups for PMA at scan, GA at birth, and a residual of the global median diffusion metric corrected for PMA and GA (see (Gondová et al., 2023)).

Metric values were then scaled between [0,1] after pooling together values across all regions and subjects within a group (PT:ses1, PT:ses2, FT). Group-specific microstructural connectivity matrices were then computed using Pearson’s correlation after concatenating individual regional microstructural fingerprints composed of the 6 diffusion metrics (4 DTI, 2 NODDI) across all subjects within the corresponding group into a single vector and considering all pairs of grey matter ROIs.

#### Functional connectivity (FC)

Based on pre-processed rs-fMRI individual data (Fitzgibbon et al., 2020), we computed median BOLD activity over labelled ROIs, applied low-pass filtering (0.1 Hz), and standardized the time-series into Z-scores. Data was smoothed (full-width at half maximum of 3.225 mm) and trimmed (first and last 50 time-points). For each subject, Pearson’s correlation was used to compute a region-based connectivity matrix from the time series of each region pair. Group-level connectivity matrices were then obtained by averaging individual matrices within each infant group (PT:ses1, PT:ses2, FT). No additional correction for confounders was included in the computation of group-wise FC. Even though the anatomically driven parcellation might not be completely adequate for delineating the functional ROIs in the developing brain (*Supp. Figure 2.3.*) we decided to keep this common framework to reduce the dimensionality of the connectomes and to allow for direct comparisons with the MC.

### Evaluation of group-wise connectivity matrices

Analyses were performed either considering all pairs of ROI connections or grouping the pairs into different subtypes: cortico-subcortical, cortico-cortical connections considering inter-hemispheric homotopic and non-homotopic, and intra-hemispheric connections. Given the importance of thalamo-cortical connectivity and the development of primary sensorimotor and visual networks during the preterm period, we further highlighted connectivity that included thalamo-cortical pairs, as well as connections involving primary cortical sensorimotor ROIs (precentral and postcentral gyri, paracentral lobule), and visual ROIs (pericalcarine cortex, lateral occipital cortex, cuneus) whose delineations were available in the current parcellation scheme.

#### Analysis of group-wise microstructural connectivity (MC)

On the MC level, we investigated the differences of the ROI connections in terms of their microstructural profile between groups (comparing PT:ses2 vs FT and PT:ses1 vs PT:ses2) and compared the distribution of the correlations between groups. As the correlation coefficients were not distributed normally in each group according to the Shapiro-Wilk test, and were considered as paired measures between groups, we used a non-parametric Wilcoxon signed-rank test to assess the differences of distributions across the group pairs. Distribution of connectivity strength between infant groups was also assessed using a robust linear regression to describe potential relationships in the patterns of MC connectivity. In this work, we employed the robust linear models with Huber’s T loss from statsmodels (v0.12.1) python package. We represented the MC connectomes as circos plots connecting ROIs. To ease the visualisation, the MC matrices were thresholded to show only the strongest 25% connection with: *i*, a common threshold across all three infant groups to uncover potential global changes of MC with age and prematurity (MC threshold r of 0.786), and *ii*, with threshold adapted to each infant group to visualise potential changes in the relative connectivity strengths between groups (adapted threshold of 0.657 for PT:ses1, 0.856 for PT:ses2, and 0.833 for FT) (presented in *Supp. Figure 4.1.*).

#### Analysis of group-wise functional connectomes (FC)

We performed similar analyses as in the case of MC to evaluate the differences in FC between the infant groups. For the creation of the circos plots from the FC connectomes, the strongest 25% connections corresponded to a common FC threshold of 0.448, and to adapted thresholds of 0.349 for PT:ses1, 0.409 for PT:ses2, and 0.537 for FT (*Supp. Figure 5.1.*).

#### Relating MC and FC modalities

The relationship between group-wise MC and FC was evaluated by robust linear regression for each infant group. The reported p-value for the slope of the described relationship was obtained by permutation testing during which the null distribution was generated by randomly shuffling the MC and FC inputs to the linear regressor. The final value was then computed as the proportion of observations more extreme than the one observed for the unshuffled inputs after 1000 random runs. The slopes were also compared using Z-scores (*Supp. Figure 6.1.*).

#### Longitudinal analysis of MC and FC modalities

We leveraged the longitudinal aspect of our dataset to evaluate potential similarities between evolution of MC and FC connectomes with age. We first computed the matrices of developmental change between PT:ses1 and PT:ses2 for MC and FC separately (referred to as ΔMC and ΔFC, respectively).

For ΔMC, as the diffusion metrics were corrected for within-group age effects before the computation of the connectome, the change between connectomic strengths with age was computed as a simple difference of absolute MC values between sessions. The direction of the change then indicated an increase or decrease of the microstructural connectivity of the given ROI connection between Ses1 and Ses2.

For ΔFC, the computation of matrices of developmental change between both timepoints was similar but included an additional step to account for the variance across the individual FC matrices within PT:ses1 and PT:ses2 groups related to the association between ROI median correlations and the infants’ PMA at the individual level. As an attempt to remedy this, we first computed group-wise connection-wise confidence intervals using the standard deviations of given connections’ connectivity strength across subjects within the considered infant group. The absolute FC differences between the 2 sessions were then weighted by the overlap of the estimated confidence interval (i.e., a high overlap led to a decreased difference). The sign of the resulting matrices of developmental change (like for ΔMC) indicates the direction (i.e., increase or decrease) of the ROI connections’ evolution with age.

The relationship between ΔMC and ΔFC in preterm infants was then evaluated by a robust linear regression applied to the components of the upper triangle of the matrices. Additionally, we compared ΔMC and ΔFC to the connectivity matrices of the opposite modality derived at two sessions (i.e., ΔMC vs FC-PT:ses1 or FC-PT:ses2, and ΔFC vs MC-PT:ses1 or MC-PT:ses2). Such analysis might allow us to assess hypotheses regarding the potential co-evolution of the MC and FC connectivity in the age-ranges of the subjects available in this study. Specifically, if the MC and FC co-evolve, their ΔMC and ΔFC networks should be highly correlated, whereas in case of stronger effect of MC on FC in the early period, we would expect ΔFC to depend on MC in the PT:ses1 group while FC at PT:ses2 would depend on ΔMC (and vice versa). To decipher if one of these three hypotheses is more relevant than the others in our PT group, we compared the regression slopes of the evaluated relationships using a Z-scores like before.

Of note, across the article, all histograms, scatter plots, and statistical comparisons include only the upper triangle of symmetric connectivity matrices. And correction for multiple comparisons refers to Benjamini-Hochberg false-discovery rate correction.

### Delineation and assessment of MC and FC networks

#### Group-wise networks

To extend our analyses beyond the direct comparisons of ROIs connection correlations, we aimed to evaluate the similarities between the inter-regional relationships to compare either the groups within each connectomic modality or between modalities. With this aim, we extracted ‘networks’ for each group and each modality separately, i.e., clusters that would regroup ROIs with similar connectivity profiles, by computing Euclidean distances from the correlation coefficients to create group-wise MC and FC distance matrices using cosine theorem (for the MC, the absolute values of the connectivity strengths were considered). We then performed hierarchical clustering with the Ward linkage to group ROIs with similar connectivity patterns. Determining the optimal number of clusters that would appropriately reflect the evolving relationships across modalities and infant groups is difficult, especially given the dynamic evolution of developing infant brain on both structural and functional level. To address this, the commonalities between hierarchical trees (dendrograms) defining MC/FC networks (either between groups for a single modality, or between modalities for a single group) were instead compared by computing mutual information (MI) across all possible cluster sizes, ranging from 2 to 75 (the maximum possible number of clusters given the number of parcels, *Supp. Figure 7.1.-2.*). MI quantifies the shared information between clustering results and approximates the overlap between network structures. Similarly to previous MC-FC comparisons, we performed permutation testing to evaluate the statistical significance of the observed MI values and account for randomness. During this procedure, cluster assignments were randomly shuffled 100 times, and the MI was recalculated to generate a null distribution of MI values for each cluster pair (2-75) in a given comparison. Observed MI values exceeding the 95^th^ percentile of the given null distribution were considered to represent meaningful, non-random clustering overlap. For the overall comparison of dendrograms, MI values for significant cluster pairs were averaged, and their standard deviations were calculated with the aim to ensure that only statistically robust overlaps contributed to the reported results, providing a more reliable measure of shared structure between networks.

#### Evolution of longitudinal MC and FC networks

Based on the idea that ROIs that participate in the same networks might show similar developmental dynamics, we further aimed to regroup ROIs based on their changing connectivity profiles, i.e., the matrices of developmental change ΔMC and ΔFC, into structural and functional ‘longitudinal networks’ in preterm infants and compare these longitudinal MC and FC-derived networks in terms of their overlap with MI. To get a proxy of longitudinal MC and FC networks, the ΔMC and ΔFC matrices were used to cluster ROIs with similar developmental connectivity modifications. As previously done for within-group connectivity matrices, we used the Ward hierarchical clustering and created all possible cluster sizes from 2 to 75 and evaluated the MI between the clustering pairs considering different combinations. Additionally, we compared the networks derived from the ΔMC and ΔFC to the networks from the respective opposite modality derived at both sessions (i.e., ΔMC vs FC-PT:ses1 or FC-PT:ses2, and ΔFC vs MC-PT:ses1 or MC-PT:ses2) to assess the potential co-evolution of the MC and FC networks (Figure 5., *Supp. Figure 7.3.*).

## Results

### Microstructural and functional relationships across grey matter

#### Evaluating microstructural connectivity (MC) in infant groups

Univariate analyses of diffusion metrics resulting from DTI and NODDI models confirmed region-specific differences between groups of infants across cortical and subcortical ROIs (*Supp. Materials: SI3. Univariate analyses of GM microstructure*).

Circos plots of the MC matrices for the three infant groups (Figure 2a, after grouping cortical ROIs into 6 lobes, and sub-cortical ROIs together, and thresholding to the 25% strongest correlation coefficients) revealed a global reinforcement of microstructural connectivity (i.e., higher correlation across ROIs) with increasing PMA (PT:ses1 vs PT:ses2/FT) across most ROI connections. In the preterm period, MC correlations were both weaker and more widespread across ROI connections within and between lobes. Among the strongest connections, some cortico-subcortical relationships observed in the preterm timepoint (e.g., subcortico-cingulate connections) were replaced by inter-hemispheric connections close to TEA. Moreover, strong negative correlations, predominately involving cortico-subcortical connections between frontal lobe and brainstem or bilateral thalamus, were observed in PT:ses2 and FT groups but not in PT:ses1 group. While challenging to attribute to a single diffusion metric, these diverging profiles are likely due to the crossing of white matter fibres in subcortical structures that give rise to opposing relationships of the related FA and ODI values at TEA (*Supp. Figure 3.2.*). This is consistent with previously described microstructural dissimilarity characterised by distinct maturational trends of diffusion metrics observed in thalamus and other subcortical ROIs compared to cortical GM (Eaton-Rosen et al., 2015; Galdi et al., 2020). To increase the comparability across groups, we considered absolute MC correlation coefficients in the subsequent analyses.

**Figure 2.**
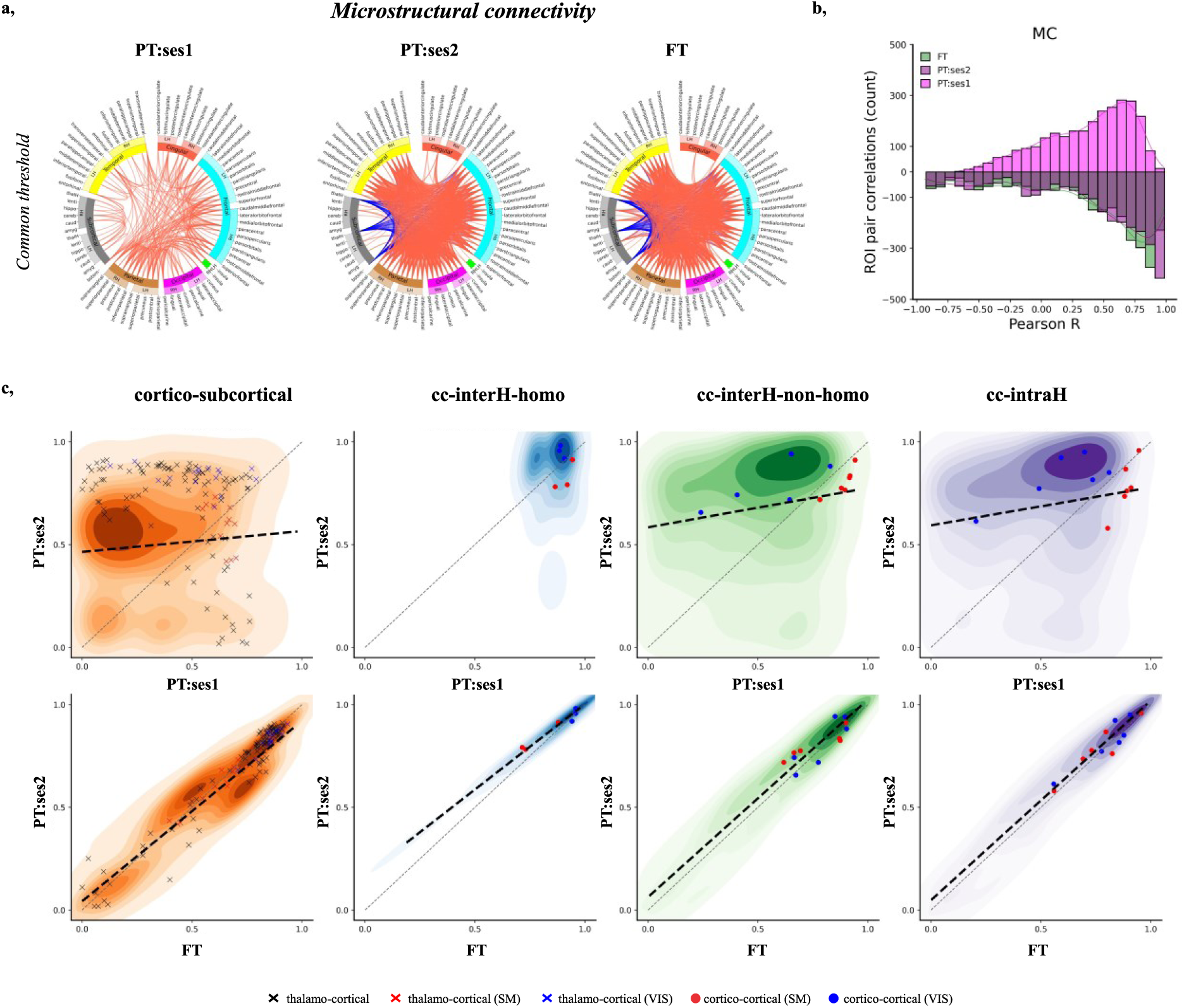
Microstructural connectome (MC) in infant groups. **a,** Circos plots representing correlation matrices and visualizing top 25 % MC connections for each infant group using a common MC threshold of 0.786. For the ease of visualization, cortical ROIs were grouped into 6 lobes (frontal in light blue, parietal in brown, temporal in yellow, occipital in pink, cingulate in red, and insular in green) and subcortical ROIs (in grey) (see *Supp. Figure 2.2.* for ROI naming conventions and assignment to lobes). Connections with positive correlations are shown in red, negative in blue. **b,** Distribution of ROI connection correlations (Pearson coefficient R) across infant groups (PT-ses1 / ses2 in light/dark magenta, FT in green) (NB: throughout the manuscript, histograms, scatter plots, and statistical comparisons include only the upper triangle of symmetric matrices). **c,** Absolute changes of MC for subsets of ROI connections between PT:ses1 vs PT:ses2 (top) and FT vs PT:ses2 (bottom). The dotted black line shows the significant* relationships determined by robust linear regression, while the grey line represents the identity relationship. Connections involving primary sensorimotor and visual regions are highlighted as scatter points. ***Legend:*** cc-interH-homo – cortico-cortical interhemispheric homotopic, cc-interH-non-homo – cortico-cortical interhemispheric nonhomotopic, cc-intraH – cortico-cortical intrahemispheric, cor.p – p value after Benjamini-Hochberg false-discovery rate correction, MC – microstructural connectivity, SM – sensorimotor, VIS – visual. *after Benjamini-Hochberg false-discovery rate correction.

Comparing ROI connection strengths between groups (Figure 2b) confirmed significant changes of MC with age, with significant differences between PT and FT infants at TEA (paired Wilcoxon test across absolute correlation values for ROI connections, corrected for multiple comparisons: PT:ses2 > PT:ses1, W=915823 p<0.001; PT:ses2 > FT, W=1521116 p<0.001. Combined with a weaker, but significant positive linear relationship between the PT:ses1 and PT:ses2 as assessed with robust linear regression, the results thus suggest an ongoing development of MC profiles in preterm neonates before TEA. As expected, much higher similarity of MC was observed between the PT:ses2 and FT groups (*Supp. Figure 4.1.*). Next, we focused on different subsets of ROI connections to evaluate potential differences in MC profiles that might reflect their different microstructural maturational patterns (Figure 2c). For cortico-subcortical connections, the MC strengths were initially mostly low (PT:ses1), and strengthened with development (PT:ses1 < PT:ses2, FT). In particular, thalamo-cortical connections showed overall high variability at all timepoints, except for those involving the primary sensorimotor (SM) and visual (VIS) areas which already showed strong connectivity at the early age. Within cortico-cortical connections, inter-hemispheric homotopic connections showed strong MC across all groups, suggesting a limited maturation effect over the studied developmental period. Other inter-and intra-hemispheric connections showed intermediate MC profiles and changes compared to cortico-subcortical and cortico-cortical homotopic connections. ROIs involved in the primary SM system showed higher MC connectivity strengths than those involved in the primary VIS system in the PT:ses1, with a comparatively lower increase in connectivity strength in the TEA timepoint.

#### Relating microstructural and functional connectivity

Estimation of FC in our work followed commonly used approach based on temporal correlation of the functional signal across the ROIs (Taymourtash et al., 2023), and the evaluation followed similar analyses as presented in the previous section for MC. The corresponding FC results are presented in the *Supp. Materials: Evaluating functional connectivity (FC)* in infant groups, as the original focus of our study was majorly MC and its relationship to functional development.

When comparing MC-FC during development across ROI connections, MC and FC showed strong positive linear relationship at an early age (PT:ses1 – slope=0.257, permuted p=0.001) which however decreased with development (PT:ses1 – slope=0.100, permuted p=0.013; FT – slope=0.046, permuted p=0.134) *(Supp. Figure 6.1a*), with statistically-different slopes between the 2 PT sessions (PT-ses1 vs PT-ses2: Z=14.02, p<0.001) and between the 2 TEA sessions (PT:ses2 vs FT: Z-score=4.29, p<0.001) (similar results were obtained when considering signed instead of absolute MC values: *Supp. Figure 6.1b*; and when analysing ROI connections with negative and positive MC values separately: *Supp. Figure 6.1c*). Considering different subsets of connections allowed us to specify these observations: positive linear MC-FC relationships were strong for all subsets in PT-ses1, whereas such relationships were observed in PT-ses2 only in connections that could be expected to be less mature at the structural and functional levels over this period (i.e., cortico-cortical inter-hemispheric non-homotopic and intra-hemispheric connections). In FT neonates, negative linear MC-FC relationships were observed for cortico-subcortical connections (Figure 3., *Supp. Figure 6.1d*).

**Figure 3.**
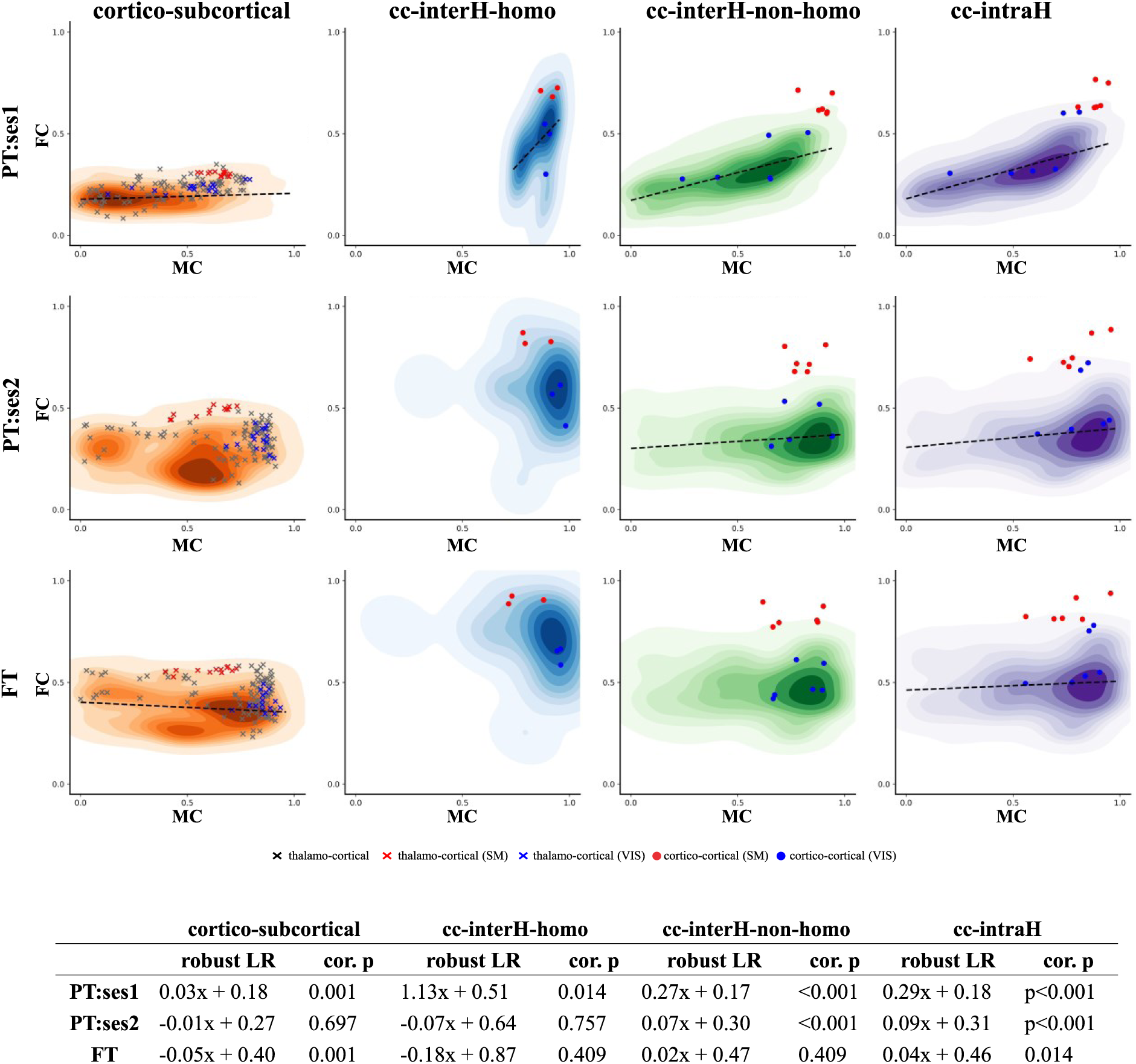
Relationship between MC and FC for different subsets of ROI connections. The table summarizes features of the robust linear relationship (LR) between the two. See Figure 2 for legend and colour codes.

#### Evaluating the longitudinal evolution of MC and FC connectomes

We further aimed to investigate the evolution of MC-FC relationships by taking advantage of the longitudinal evaluations of PT infants to compute the matrices of developmental change between PT:ses1 and PT:ses2 separately for MC (considering absolute values at each age) and FC (referred to as ΔMC and ΔFC, respectively) (Figure 4a). Both increases and decreases of the connection strengths from the preterm period to TEA were observed for both modalities. As seen from group-wise comparisons, most connections increased in strength for ΔMC. Among others, the most pronounced decreases in connections involved the cingulate cortex and some of the subcortical ROIs (e.g., cerebellum and amygdala). For ΔFC, connections involving subcortical ROIs globally increased in strength while several cortico-cortical connections decreased in strength in a more pronounced way than for ΔMC. Inter-hemispheric connections involving homotopic cortical ROIs showed different profiles compared to other cortico-cortical connections (inter-hemispheric non-homotopic and intra-hemispheric connections) which were merged in the subsequent analyses given their overall similarity.

**Figure 4.**
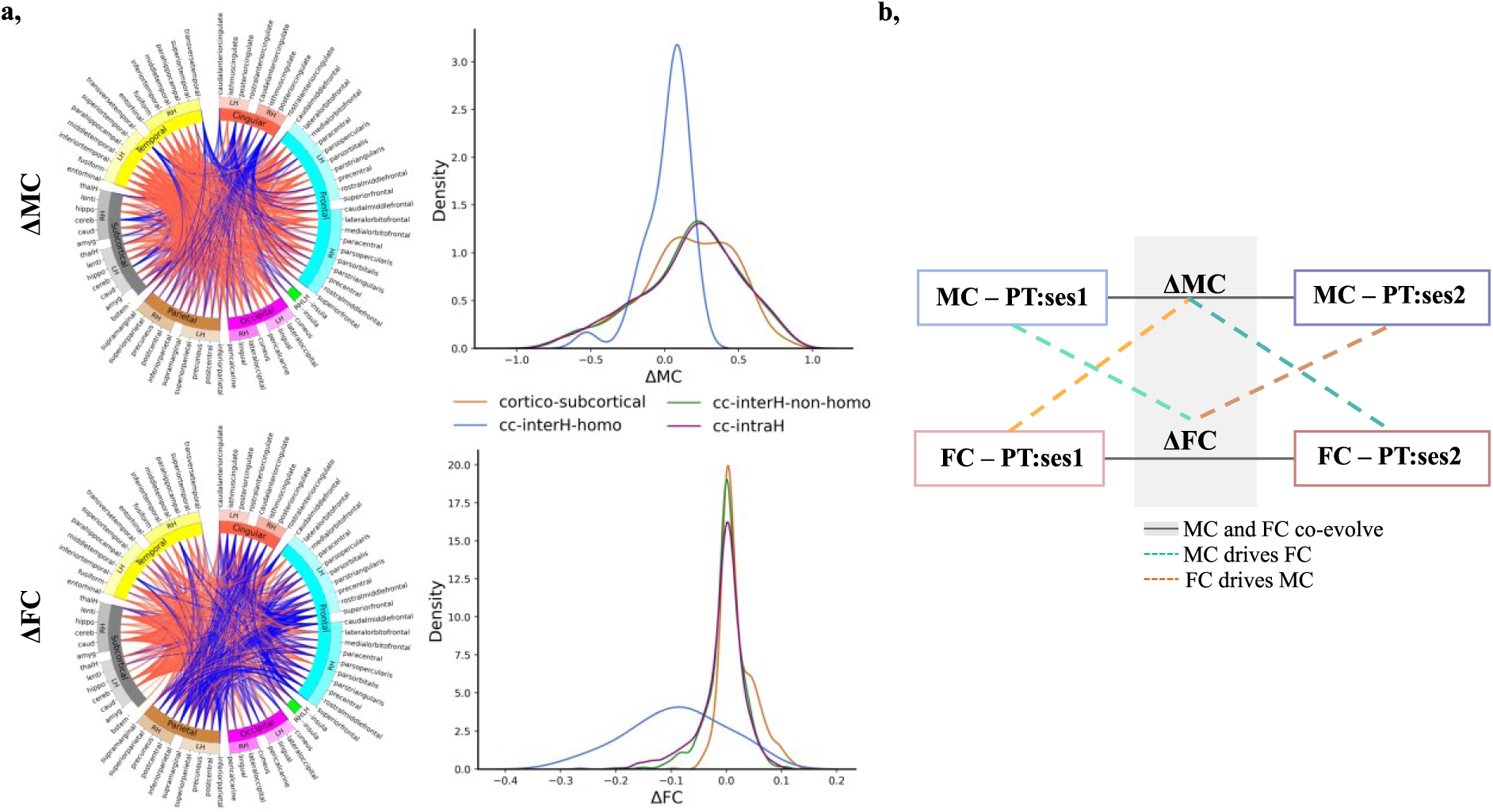
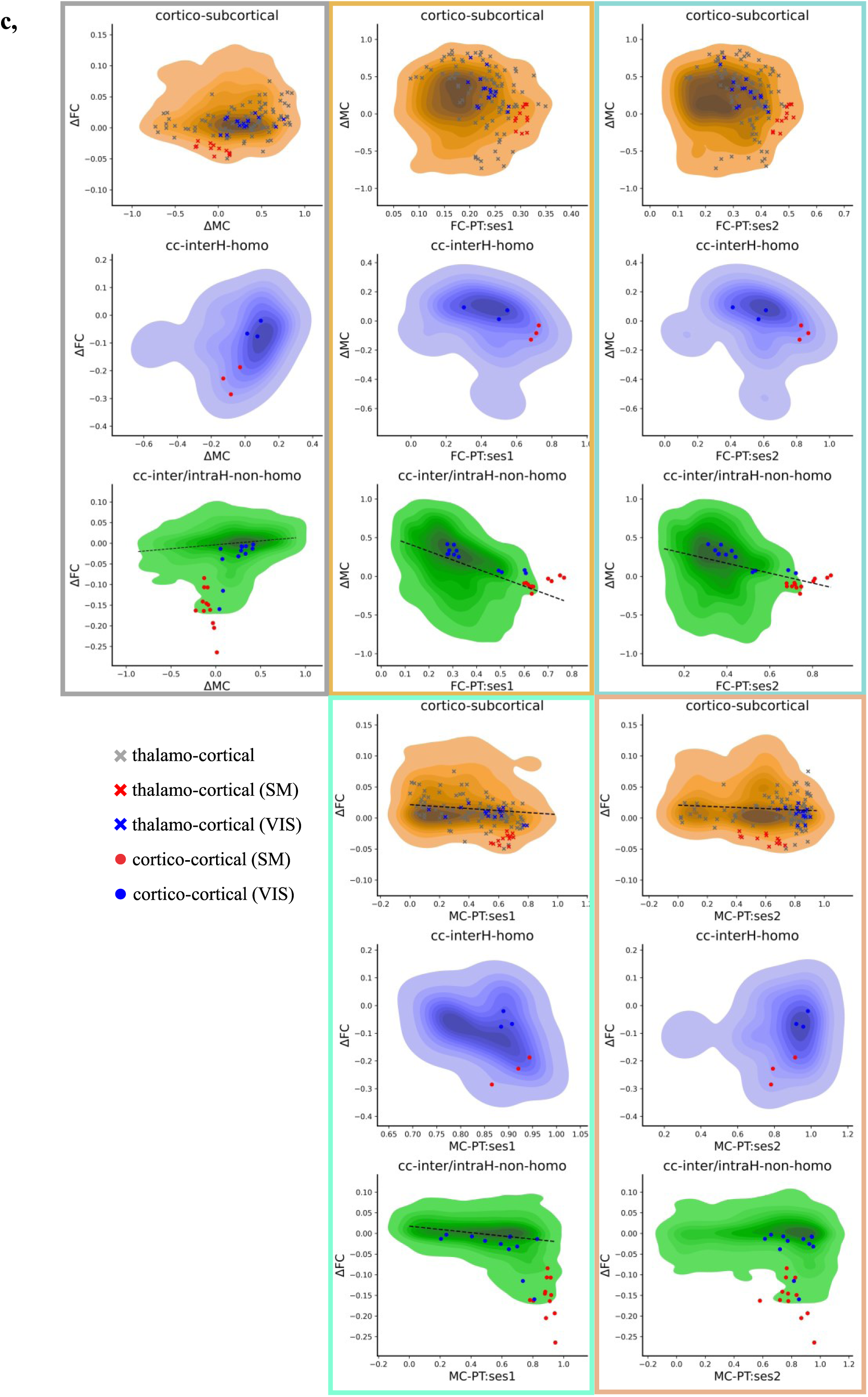
**a,** Longitudinal evolution of MC (top left) and FC (bottom left) in preterm infants. Circos plots visualize top 25 % connections for ΔMC and ΔFC modalities in PT infants (respective thresholds: 0.458 and 0.037). Connections with increasing strength of relationships with age are shown in red, decreasing in blue. Density plots show the distribution of changes in connectivity strength with age for different subsets of connections for the two modalities (right). **b,** Figure summarizing 3 possible hypotheses on the developing causal relationships between microstructural and functional connectivity: we colour coded the subplots by hypothesis tested by the given comparison: MC and FC co-evolve (grey), MC drives FC (cyan, dark cyan), and FC drives MC (orange, dark orange). **c,** Directionality of MC-FC relationships for different connection subsets (colour-coded by the three hypotheses). See Figure 1 and *Supp. Figure 2.2.* for legend and colour codes.

To further investigate the potential directionality of dependence between the developing MC and FC in preterm infants, we hypothesized three possible MC-FC relationships (Figure 4b): 1) if FC and MC co-evolve, ΔFC and ΔMC should be strongly correlated; 2) if FC relies on MC, MC-ses1 should drive ΔFC while ΔMC should drive FC-ses2; 3) conversely, if MC relies on FC, FC-ses1 should drive ΔMC while ΔFC should drive MC-ses2. The hypotheses and results for all ROI connections are summarized in *Supp. Figure 6.2b,c*, while here we focused on the subsets of connections (Figure 4c).

For cortico-subcortical connections, significant negative linear relationships were observed between ΔFC and MC-ses1 but also between ΔFC and MC-ses2. Among cortico-cortical connections, inter-hemispheric homotopic connections showed no significant associations across the comparisons, which could be related to their relatively mature state compared to other cortico-cortical connections during the considered age range or by methodological limitations due to reduced number of connections compared to other subsets. For the other cortico-cortical connections (inter-hemispheric non-homotopic and intra-hemispheric), ΔFC and ΔMC were positively related, while ΔMC and FC-ses1 as well as ΔFC and MC-ses1 showed negative relationships. This latter result suggests that there exist synchronized changes in MC and FC, and also lower changes in one connection modality if the other already showed high connectivity strength during the preterm period. Surprisingly, negative relationships were also observed between ΔMC and FC-ses2, suggesting a reverse pattern (e.g., larger developmental increases in MC “leading” to lower FC at ses2).

### Network-based comparisons of MC and FC development

To extend the direct comparisons of connectivity strength across ROI connections between infant groups, we further used the connectivity matrices to define ‘microstructural networks’ for MC and ‘functional networks’ for FC modality using Ward clustering for each infant group. The resulting dendrograms, as well as examples of clustering for selected cluster numbers are presented in *Supp. Figures 7.1. and 7.2*.

In agreement with previous observations, the comparison of clustering results using mutual information (MI) between hierarchical trees across infant groups for either MC or FC showed higher, although imperfect, overlap between PT:ses2 and FT subjects than between the 2 PT sessions (Figure 5a), while differences between PT:ses2 and FT subjects supported potential effects of prematurity on microstructural connectivity across ROI connections. Interestingly, the network overlap was very similar in FC and MC modalities, whereas a stronger positive linear relationship between PT:ses1 and PT:ses2 groups in terms of FC than of MC was previously observed across ROI connections. This suggested that network structures observed at TEA are already in place in the preterm period to a certain extent for both FC and MC.

**Figure 5.**
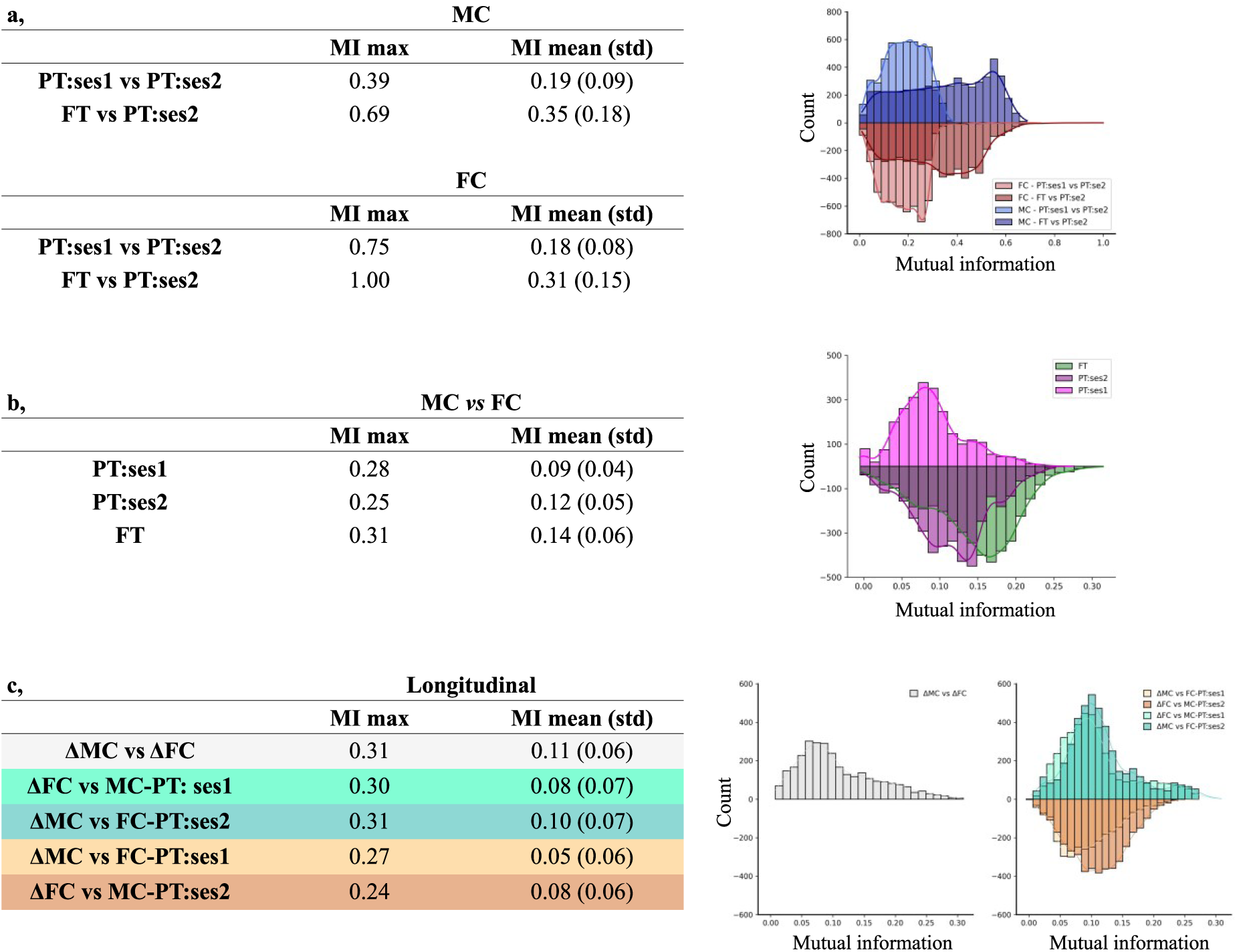
Network-based analyses. **a,** Table and histogram summarising the distribution of mutual information (MI) of clustering results within MC (blue) and FC (red) modalities between groups (PT:ses1 vs PT:ses2 in light colour, FT vs PT:ses2 in deep colour). **b,** Results for the network overlap comparisons between MC and FC modalities. Distribution of MI values within PT:ses1 group in light magenta, PT:ses2 dark magenta, and FT in green. **c,** Mutual information between clustering results derived from longitudinal ΔMC and ΔFC matrices in preterm infants as well as comparison to networks derived from the opposite modality at ses1 or ses2. As in Figure 4, we colour coded the columns by hypothesis tested by the given comparison: MC and FC co-evolve (grey), MC drives FC (cyan), and FC drives MC (orange).

Using the clustering-based approach, we also evaluated the similarity between MC and FC at the network level in each group using the clustering approach. A trend of low but progressively increasing mutual information was observed between MC- and FC-derived networks (i.e., lower mean values in PT:ses1 than in FT, Figure 5.b) which contrasted with direct ROI connection comparisons where the MC-FC linear relationships tended to globally decrease with development. This might suggest a potential emergence of the shared underlying network structures between MC and FC throughout development.

Finally, as in previous sections, we performed analysis of hierarchical clustering separately on ΔMC and ΔFC to characterise longitudinal networks. Derived ΔMC and ΔFC dendrograms and examples of clustering results for selected number of clusters are presented in *Supp. Figure 7.3.,* displaying some visual similarities but also some dissimilarities between ΔMC and ΔFC. The mutual information between clustering results from longitudinal ΔMC and ΔFC was further compared to the networks at each PT session to test the MC-FC relationship hypotheses proposed in Figure 4b. Evaluation of MI between ΔMC and ΔFC derived networks suggested significant overlap, which tended to be higher than for all the other comparisons between each longitudinal network and the opposite modality at a given session (Figure 5c). The lower MI were observed for network comparisons of ΔFC vs MC-ses2 and FC-ses1 vs ΔMC, suggesting a lower dependence of MC on FC in the network space than the reverse (Figure 5c). Nevertheless, results across the comparisons were fairly similar, making it difficult to confidently ascertain which of the three hypotheses about MC-FC co-evolution is the most probable on the network level.

## Discussion

This study investigated early postnatal changes in *microstructural* and *functional* brain connectivity, with a focus on their developmental trajectories during the preterm period. We observed a global strengthening of absolute connectivity across cortical and subcortical regions. However, these changes varied significantly across connections, consistent with the expected stages of connection maturation, and were systematically impacted by prematurity.

Our findings imply that both microstructural and functional grey-matter connectivity not only strengthen with age but also appear to evolve in relation to each other. This is highlighted by significant positive linear relationships between MC and FC in the early PT session, suggesting a potentially tightly coupled developmental trajectory between these modalities. Later progressive decrease in direct positive MC-FC relationship with age may reflect a dynamic shift in how the two modalities relate to one another, potentially reflecting a transition toward greater functional specialization and complexity as the brain matures.

Interestingly, while direct MC-FC relationships diminished over time, our findings suggest that the overlap between derived MC and FC networks increased. This might rely on a progressive refinement and potential convergence of network structures between the two modalities. Such convergence may reflect the establishment of efficient interactions across the neural networks to support maturational processing and integration during this critical period of brain development.

### Early maturation of grey matter connectivity

#### Evaluating early microstructural connectivity (MC) with multi-shell dMRI

By analysing microstructural features, our study provides new insights into early neurodevelopment through an alternative approach that evaluates microstructural relationships (i.e. connectivity) across GM regions in relation to their functional synchronicity in the temporal domain. Covariation of GM features during the first two years after birth have been studied in the past, primarily relying on single morphometric markers (Fan et al., 2011; Geng et al., 2016; Nie et al., 2014). While informative, such approaches offer only an indirect view of the underlying microstructural processes occurring during the complex period of early brain development (including dendritic arborization and the growth of axonal extensions, synaptogenesis and subsequent pruning, myelination of intracortical fibres, proliferation of glial cells, which together lead to a dramatic decrease in water content and an increase in tissue density) (Bystron et al., 2008). Additionally, observations based on single descriptors are potentially biased by specific spatial and temporal developmental patterns of changes of the given descriptor (Gilmore et al., 2012; Lyall et al., 2015; Nie et al., 2014; Seidlitz et al., 2018). We thus aimed to achieve a more comprehensive view of the inter-regional GM connectivity by leveraging the complementary microstructural information provided by DTI- and NODDI-derived metrics.

Both dMRI models present different trade-offs between complexity, biological plausibility and limitations, such as those due to the DTI model inability to accurately represent microstructure in regions with complex voxel properties (Batalle et al., 2019) or the potential suboptimal estimation of NODDI parameters in infant GM due to its initial optimisation for adult WM (Guerrero et al., 2019). However, both models have been widely used in developmental studies because of their relevance as GM microstructure descriptors (Ball et al., 2013; Eaton-Rosen et al., 2017; Smyser et al., 2016). To maintain consistency with the literature, we opted for similar settings as previous dMRI studies of cortical maturation for NODDI estimation (Batalle et al., 2019; Fenchel et al., 2020), and extracted metrics over the cortical surface (Lebenberg et al., 2019) while considering median values over ROIs (Gondová et al., 2023) for a reliable evaluation of GM microstructure confirmed by the observation of expected diffusion metric changes with age (*Supp. Figure 3.1. & 3.2.*).

Combining multiple morphological and microstructural descriptors of cortical development into a multi-parametric approach has been previously proposed in FT infants to identify modules and similarity networks consistent with known cytoarchitecturally defined brain areas and functional systems (Fenchel et al., 2020; Galdi et al., 2020). Nevertheless, morphological features from anatomical MRI (e.g. local cortical surface, thickness, and folding) remain less specific markers of maturation synchronization than microstructural features employed in our study with dMRI. Due to the limited descriptor set (6 microstructural metrics), we focused on group-wise MC estimation that included corrections for confounding factors (i.e. GA at birth, PMA at scan). Extracted matrices remained globally consistent across groups, suggesting reliability of our approach. Future work could potentially benefit from increasing the feature sets by incorporating additional microstructural descriptors including diffusion kurtosis metrics (Jelescu et al., 2015) for computation of individual matrices.

In terms of MC, the observed global reinforcement of absolute connectivity strength across regions with increasing age agrees with previous descriptions of complex age-related changes of microstructural similarity in FT and PT infants at TEA, including increases across occipital, parietal and temporal areas, and decreases in limbic and cingulate regions (Fenchel et al., 2020; Galdi et al., 2020). Observations in frontal regions differed across studies, perhaps because of diffuse alterations of brain maturation in the subjects born preterm (Ball et al., 2017). The age-related increase in absolute connectivity observed between thalamus/brainstem and cortical regions is also consistent with rapid developmental changes to afferent and efferent deep GM connectivity reported in the preterm period (Batalle et al., 2017).

#### Evaluating functional connectivity (FC) with rs-fMRI

Several studies have highlighted the organisation and evolution of FC and related networks during early brain development (Dall’Orso et al., 2022; Eyre et al., 2021; Taymourtash et al., 2023; Williams et al., 2023). Although data used in our study were processed using an optimised pipeline designed to minimize the effects of motion, residual difference between the groups were observed. Despite the differences in data quality across the three infant groups (Supp. Table 2.1), our unimodal FC analyses (presented in the *Supp. Materials: Evaluating functional connectivity (FC) in infant groups*) revealed coherent profiles, similar to those described in previous studies (e.g., stronger connectivity between homotopic inter-hemispheric regions than between other regions (Taymourtash et al., 2023; Williams et al., 2023)). While we noted effects of PMA at scan across individual FC matrices, we did not correct for it when computing group-wise matrices due to difficulties with implementation of appropriate corrections at the level of individual connections. Nevertheless, we attempted to mitigate these effects when deriving ‘weighted’ longitudinal matrices between the preterm period and TEA (see *Materials and methods: Longitudinal analysis of MC and FC modalities*). Similarly to our MC results and previous FC studies (Taymourtash et al., 2023; Thomason et al., 2015), we observed a global FC reinforcement of connectivity strengths with age, particularly for cortico-subcortical but also cortico-cortical connections. Nevertheless, the detection of early cortico-subcortical FC at the first timepoint might be limited by age-dependent effects on signal-to-noise ratio (SNR) (Denisova, 2019) that may notably impact the subcortical regions (Maugeri et al., 2018; Risk et al., 2021). While it is interesting to note that FC between the two PT sessions showed a higher correlation than MC, interpretation in terms of maturation progression would require comparison to MC and FC mature states to disentangle whether functional connections are established earlier than microstructural ones, or if the refinement of FC relationships occurs after the studied period.

#### Relationships between microstructural and functional connectivity in the late preterm period

To provide a common framework for the comparisons of MC and FC modalities, we used a set of 31 bilateral cortical and 13 subcortical ROIs, derived from anatomical parcellations optimized for neonates. Nevertheless, the anatomically driven parcellation likely introduces bias to resulting connectivity estimates, particularly in the case of FC in which anatomically defined ROIs might not accurately represent individual functional regions especially in the context of dynamic development (Smith & Beckmann, 2017; *Supp. Figure 2.3.*). Additionally, although the parcellation quality was satisfactory for all subjects and timepoints, it was lower for the earlier scans (i.e., when the infant’s age moved away from TEA ‒ age used to define the cortical parcellation atlas) (*Supp. Figure 2.2.*). The observed errors occurred mostly in the regions where anatomical landmarks, such as secondary and tertiary sulci, were not yet present to guide the delineations. We extracted median ROI descriptors in both MC and FC modalities to partially mitigate these parcellation errors. Future work could use different parcellation schemes (e.g., random parcellation (Fenchel et al., 2020; Gondová et al., 2023)) to confirm our findings.

Notably, while there were significant positive linear relationships between MC and FC in the preterm period, coupling decreased with development at the whole brain level. Comparison between developmental matrices across all connections in the preterm period (ΔMC and ΔFC) suggested a clear trend of positive relationship between coinciding changes of MC and FC, as well as a lower change in one modality if the other already showed high connectivity strength in the early period. The initially significant positive relationship in the preterm period is followed by a progressive but coordinated decoupling of MC and FC with maturation. This contrasts with previous studies linking structure and function through white matter connectivity (Grayson et al., 2014; Hagmann et al., 2010; Zhang et al., 2022) which described increasing coupling in PT infants (van den Heuvel et al., 2015) and synchrony between the maturation of grey matter regions and underlying white matter tracts (Friedrichs-Maeder et al., 2017; Smyser et al., 2016). The approach we proposed with MC could thus provide an alternative and complementary view for investigations into structure-function relationships even in absence of identified structural links (i.e., structural connections identified with diffusion MRI and tractography).

#### Different MC-FC relationships in connection subsets based on expected maturational stage

Any evaluations of MC-FC relationships are intrinsically influenced by the considered developmental period relative to the maturation stage of the given connections and networks. Thus, whole-brain evaluations are likely obscured by the asynchrony in maturation across brain areas observed at the level of GM microstructure (Fukutomi et al., 2018; Lebenberg et al., 2019) and FC (Cao et al., 2017; Eyre et al., 2021; Larivière et al., 2020). To address this and perform more targeted evaluation of MC-FC relationships, we considered subsets of connections (i.e. cortico-subcortical vs cortico-cortical connections, with distinctions between intra-and interhemispheric ones, the latter being split into homotopic vs non-homotopic subsets) that were previously shown to differ based on previous studies of functional (van den Heuvel et al., 2015) and white matter structural connectivity (Kostović et al., 2019; Kulikova et al., 2015; Takahashi et al., 2012; Vasung et al., 2017; Wilson et al., 2021, 2023).

Across cortico-cortical connections, inter-hemispheric homotopic ones showed similar profiles of strong MC between the two PT sessions, indicating limited maturation effects over the studied period, while the other connections showed more varied MC changes. Qualitatively similar trends were observed for FC, with inter-hemispheric homotopic connections being generally similar between sessions. Cortico-subcortical connections seemed to display greater heterogeneity in age-related changes in both MC and FC strength with age, suggesting a need for more specific analysis across different deep GM structures (but this was out of the scope of this first study).

Cortico-cortical connections involving primary sensorimotor (SM) and visual (VIS) ROIs both showed strong MC at PT:ses1 with minimal developmental changes across the studied age range, in line with reports of early maturation of primary sensory areas (Ball et al., 2013; Lebenberg et al., 2019). SM regions exhibited slightly higher initial MC and a smaller increase by TEA compared to VIS regions, that might indicate earlier maturation of sensorimotor functions. Similar to MC, strong FC was observed for SM connections with minimal developmental change over the studied period, consistent with previous observations of functional organisation of primary sensory networks by TEA (Eyre et al., 2021; Dall’Orso et al., 2022). This was to a lesser degree similar for VIS connections, in line with previous findings suggesting the early presence of sensory cortical hubs with a later transition to the visual system (Fransson et al., 2009; van den Heuvel et al., 2015).

Comparing the two MC and FC modalities within each group (Figure 3.), different connection subsets revealed diverse patterns of MC-FC relationships with age. Cortico-subcortical connections exhibited negative relationships at TEA, while inter-hemispheric homotopic and non-homotopic connections showed decreasing coupling, and intra-hemispheric connections maintained significant positive relationships across all age timepoints. This observation might be coherent with the developmental sequence described in terms of white matter connectivity and FC across different connection subsets, with cortico-subcortical connections being the most mature over the perinatal period, followed by inter-hemispheric homotopic cortico-cortical connections, non-homotopic inter-hemispheric connections, then the remaining intra-hemispheric cortico-cortical connections. Progressive loss of MC-FC coupling could then reflect the connectivity maturation on the microstructural level which might underlie progressive functional specialisation and diversification of FC (Allievi et al., 2016; Dall’Orso, 2022). However, if the interpretation of progressive functional specialization is correct, the decoupling between MC and FC likely stems from the measure to which FC matrices derived from a given set of ROIs capture the underlying biological functional connectivity changes with age. Thus, while the loss of positive relationships might indicate developmental changes that occur during the studied age range between the two modalities, distinguishing between biological and ‘second-order’ methodological artefacts is challenging. Future work would benefit from adapting parcellation schemes to better reflect functional (and microstructural) specialization with age or attempting the analysis of spatial distribution of developmental changes across cortical and subcortical structures in a parcellation-independent manner.

Furthermore, the observation of negative relationships between MC and FC in potentially the most mature cortico-subcortical connections at TEA suggests a complex structure-function relationship in mature systems that warrants further exploration. As we suspected that negative MC values at later ages (resulting from microstructural differences between cortical and subcortical ROIs) might complicate these comparisons, we also analysed the MC-FC relationships separately for positive and negative MC, confirming different characteristics for cortico-subcortical vs cortico-cortical connectivity (*Supp. Figure 6.1d*).

Future MC analyses might be reserved to the cortico-cortical assessment, while targeted white-matter structural connectivity evaluations might be more appropriate to consider cortico-subcortical connectivity (Neumane et al., 2022).

While our goal was to explore the directionality of MC and FC co-development based on matrices of developmental change in PT infants (ΔMC and ΔFC), results did not allow us to distinctly differentiate between the three possible hypotheses: FC-MC co-evolution; MC relying on FC; FC relying on MC. The relative limited sample size in our study, due to the scarcity of multimodal and longitudinal data from the preterm population, may partly explain our inconclusive results. Additionally, variability in the birth-to-scan and 1st-to-2nd scan delays in the PT group could also affect group comparisons (*Supp. Figure 1.1.*). While reassigning or excluding subjects to create groups with more homogenous scan ages could reduce variability, it would further reduce our sample size. We chose to retain as many subjects as possible, controlling for age at scan as a linear covariate. Nevertheless we acknowledge that such corrections may not fully capture the complex, nonlinear developmental trajectories during this period. Future studies could address age variability more robustly by modelling age as a continuous variable, shifting to individual-level connectivity estimates which would allow to compare the rates of change across different connections in a more comprehensive manner, and expanding the longitudinal dataset to enhance the robustness of our evaluation. Until then, the observation of significant relationships in the latest developing inter/intra-hemispheric non-homotopic subsets but not in the others might be suggestive of a concurrence of bi-directional MC-FC changes in developing connections. Further targeted investigation focusing on selected networks with well characterised maturational sequences, such as primary sensorimotor and visual networks may help to further elucidate the developing interactions between MC and FC in the absence of larger longitudinal cohorts, although their limited developmental changes across the period studied might make the assessments challenging.

#### Impact of prematurity on microstructural and functional connectivity

Additionally, as previous studies have reported that PT infants show spatially heterogenous alterations in GM microstructure (Batalle et al., 2019; Dimitrova et al., 2021; Eaton-Rosen et al., 2015, 2017; Ouyang, Jeon, et al., 2019; Smyser et al., 2016) and altered inter-regional similarity (Galdi et al., 2020), we also investigated the impact of prematurity on MC and FC at TEA. We observed different relative patterns of MC between the two groups (PT:ses2 vs FT), affecting diverse ROI connections, including the bilateral thalamus and hippocampus, as well as widespread intra-and inter-hemispheric connections across the cortex. Interestingly, PT infants showed globally higher MC strengths at TEA compared to FT infants, particularly in inter-hemispheric homotopic cortical connections. This may suggest a more mature profile in PT infants for these connections, potentially reflecting accelerated maturation driven by the earlier onset of experience-dependent developmental mechanisms in the extra-uterine environment.

In terms of FC, we observed a significant effect of prematurity characterised by a global decrease in FC connectivity strength, consistently with previous studies (Ball et al., 2016; Brenner et al., 2021; Chiarelli et al., 2021; Eyre et al., 2021; Keunen et al., 2017; Scheinost et al., 2016; Smyser et al., 2010). This suggests a diffuse and complex effect of prematurity on FC, rather than more focal effects on intrinsic brain network connectivity.

When directly comparing MC and FC, differences between PT infants at TEA and FT neonates were observed only in cortico-subcortical and inter-hemispheric non-homotopic connections. These findings suggest that the impact of prematurity on MC-FC relationships may depend on the maturation stage and specific dynamics of connection subtypes, which could be influenced by the timing of the premature birth among other complex factors, including environment. In our study, PT and FT infants did not differ in socio-economic status, as measured by the UK Index of Multiple Deprivation (IMD) (*Supp. Table 2.2.*) and we considered the environmental impact on brain connectivity to be limited at the time of MRI in our study. However, environmental factors are known to influence neurodevelopment in premature (and full-term) infants (Benavente-Fernández et al., 2019; Vanes et al., 2023) and likely play a broader role in brain development through complex interactions. While our study focused on differences linked to prematurity, future work could examine how socio-economic factors affect maturation and modulate the effects of prematurity. As early life presents a sensitive window during which brain disruptions may disproportionately affect later outcomes, the MC and FC modifications observed in our work could predispose to altered trajectories and dynamics of brain network integration and specialisation with long-term implications for neurodevelopmental outcomes. Future research should explore specific effects of perinatal insults and their timing on MC-FC dynamics across maturational stages to inform more personalised intervention strategies that could optimize brain network development during critical periods.

While our study focused on the preterm period in preterm infants, prematurity-related alterations to microstructural and functional modalities make it difficult to dissociate maturation from prematurity effects. Since little is known about the developmental relationships between the structure and function of emerging neuronal networks, future longitudinal research in typical populations might be warranted to better understand these physiological dynamics and their differences in pathology.

Previous studies have further highlighted the significant role of postnatal experiences in brain network maturation. While we attempted to control for GA at birth and PMA at MRI scan in each group, the time between birth and MRI (days *ex-utero*) could still impact MC, FC, and their assessed relationship. While it would be difficult to fully assess these effects due to our limited sample size compared with GA and PMA variability, future work might investigate the effects of days *ex-utero* to better distinguish between environmental and intrinsic developmental influences.

### Network approach: sub-setting links with similar connectivity profiles and developmental trajectories

To expand our analyses beyond direct comparisons of MC and FC strengths, we aimed to assess the potential overlap of the spatial organisation of networks across infant groups and modalities. This involved defining networks based on connectivity profiles across different regions. To do so, we used hierarchical Ward clustering due to its ability to capture potential hierarchical structure of regional relationships and to retain interrelations at different levels of description in the resulting dendrograms. However, this method has limitations, among which the irreversibility of the cluster assignment that makes the resulting clusters sensitive to local effects and errors that might be propagated through dendrograms (Moreno-Dominguez et al., 2014). Despite this, Ward clustering was previously shown to perform well compared to alternative methods, even for a large number of clusters, in both simulated and real rs-fMRI data (Thirion et al., 2014). Nevertheless, given the potential spatial overlap between brain networks, future research could explore other clustering methods such as overlapping communities (de Reus et al., 2014) or those accommodating varying spatial network configurations (Bijsterbosch et al., 2018). Additionally, more granular analyses with larger number of smaller ROIs could be beneficial, especially in the case of FC based on random or functionally based parcellations.

Determining an appropriate number of clusters also posed challenges, especially because the network specialization might differ throughout development and between MC and FC modalities (Allievi et al., 2016; Dall’Orso et al., 2022). To address this, we compared all possible cluster number pairs to approximate general overlap across different network solutions using the mutual information. Although this approach provided only a broad measure of network (di)similarity, visual inspection of resulting hierarchies suggested informativeness within the clustering solutions, with some expected patterns such as regions within primary sensorimotor networks tending to belong to the same clusters across solutions.

Such a network-based approach might thus offer alternative and complementary information to previous connection subset-based descriptions. Simplifying the heterogeneity across connections by clustering regions with similar connectivity profiles (for group-specific clustering) or similar developmental trajectories (for longitudinal networks between PT:Ses1 and PT:Ses2) to derive connectivity clusters, i.e., ‘networks’, might lead to more robust comparisons of MC and FC connectivities.

At the network level, we observed significant but not extensive overlap between MC-derived networks in preterms (PT:ses1 vs PT:ses2) and at TEA (PT-ses2 vs FT), suggesting both convergences and divergences over the preterm and term periods. Nevertheless, some patterns of microstructural connectivity reflecting potential network properties appear to be established during the preterm period and further refined with development. Similarly, FC-derived networks showed some overlap between the two PT sessions, suggesting that some functional properties are discernible in the preterm period and continue to develop as suggested by previous studies (Doria et al., 2010; Smyser et al., 2010; van den Heuvel et al., 2015). Besides, the observation that the network overlap remains consistent between the two PT sessions for both MC and FC modalities contrasts with the direct connectivity comparisons indicating higher similarity of FC than MC, further underscoring the value of the network-level analysis to describe inter-regional patterns that might be inaccessible in the whole-brain analysis due to heterogenous maturation of different connection subtypes.

Interestingly, while direct connectivity strength comparisons indicated disappearance of MC-FC linear relationships with maturation, comparisons of extracted networks revealed an opposite trend of increasing network overlap with age. This might imply a shared underlying network organization between MC and FC established early on in the preterm period, similarly to later ages (Geng et al., 2016), that progressively aligns MC and FC as networks refine on both microstructural and functional levels. Network-based comparisons derived from developmental changes (ΔMC and ΔFC) reached similar overlaps, further suggesting network commonalities between MC and FC modalities. As for direct MC-FC comparisons, typical organisation of MC-FC network patterns seemed altered by prematurity indicated by a lower overlap in PT:ses2 infants than in FT neonates.

Regarding the question of MC-FC co-evolution and directionality, we observed generally lower network-based overlap for the comparisons ΔMC vs FC-PT:ses1 and ΔFC vs MC-PT:ses2 that weakens the hypothesis of dependence of MC on FC during the preterm period. Instead, the two alternative hypotheses (MC-FC co-evolution or dependence of FC on MC) might be better supported by such network-based observations. Nevertheless, given the complex and bi-directional influences between microstructural and functional development underlying brain maturation in this period (Cadwell et al., 2019), further investigations focusing on specific systems/networks might be necessary to reduce analytical noise and clarify the complex picture of MC-FC relationships. Additionally, previous studies suggested that the relationship between white-matter structural and functional connectivity might diminish as structure stabilizes into a more permanent foundation for the adaptations of functional connectome to new demands and environment (Baum et al., 2020; Ciarrusta et al., 2022; Yeo et al., 2011). Extending our evaluations of MC-FC relationships to later developmental periods, when the interplay between microstructure and function may evolve differently, might be useful to complement our observations.

## Conclusion

In the present study, we explored the complex nature of grey matter connectivity during early brain development through comparisons of microstructural and functional brain features. Focusing on 45 preterm infants scanned longitudinally, we observed a global reinforcement of absolute MC and FC strength with age, characterise by strong dependence on different connection subsets and network maturational dynamics. MC and FC are positively related during the preterm period but this linear relationship decreases with development, while overlaps between MC-and FC-derived networks increase with age, suggesting a progressive convergence toward a shared network structure. These findings highlight the intricate interplay between microstructural and functional properties and will hopefully lead to future studies into how their co-evolution may play critical role in shaping neurodevelopmental trajectories and their disruption impact long-term outcomes.

Prematurity had a diffuse and heterogeneous effect on both MC and FC, with significant reductions in connectivity observed in preterm infants compared to their full-term counterparts. These disruptions underscore the need for further research to investigate how specific MC-FC alterations relate to the degree of prematurity and how they influence later neurodevelopmental outcomes. In the future, examining individual-level variations in MC-FC relationships and their progression through later developmental stages may help delineate atypical trajectories in vulnerable preterm populations. Such efforts could enhance our understanding of neurodevelopmental disorders and inform targeted interventions for preterm infants in order to improve their functional outcomes and quality of life.

The increasing overlap between MC and FC networks with age also emphasizes the potential utility of MC as a complementary descriptor for characterizing brain network maturation. While the biological significance of MC in synchronized maturation across brain regions warrants further investigation, future studies comparing MC with white matter structural connectivity and exploring its relationships to intrinsic developmental triggers (e.g. conserved genetic or evolutionary patterns), extrinsic environmental influences and subsequent behavioural acquisitions may provide additional insights.

## Data availability

The employed dataset is available from: http://www.developingconnectome.org/data-release/third-data-release/.

## Ethics approval / patient consent statement

The dHCP project received UK NHS research ethics committee approval (14/LO/1169, IRAS 138070), and written informed consent was obtained from the parents of all participant infants.

## References

Adamson, C. L., Alexander, B., Ball, G., Beare, R., Cheong, J. L. Y., Spittle, A. J., Doyle, L. W., Anderson, P. J., Seal, M. L., & Thompson, D. K. (2020). Parcellation of the neonatal cortex using Surface-based Melbourne Children’s Regional Infant Brain atlases (M-CRIB-S). Scientific Reports, 10(1), 1–11. 10.1038/s41598-020-61326-2

Alexander-Bloch, A., Giedd, J. N., & Bullmore, E. (2013). Imaging structural co-variance between human brain regions. Nature Reviews Neuroscience, 14(5), 322–336. 10.1038/nrn3465

Allievi, A. G., Arichi, T., Tusor, N., Kimpton, J., Arulkumaran, S., Counsell, S. J., Edwards, A. D., & Burdet, E. (2016). Maturation of Sensori-Motor Functional Responses in the Preterm Brain. Cerebral Cortex, 26(1), 402–413. 10.1093/cercor/bhv203

Ball, G., Aljabar, P., Arichi, T., Tusor, N., Cox, D., Merchant, N., Nongena, P., Hajnal, J. V., Edwards, A. D., & Counsell, S. J. (2016). Machine-learning to characterise neonatal functional connectivity in the preterm brain. NeuroImage, 124, 267–275. 10.1016/j.neuroimage.2015.08.055

Ball, G., Aljabar, P., Nongena, P., Kennea, N., Gonzalez-Cinca, N., Falconer, S., Chew, A. T. M., Harper, N., Wurie, J., Rutherford, M. A., Counsell, S. J., & Edwards, A. D. (2017). Multimodal image analysis of clinical influences on preterm brain development. Annals of Neurology, 82(2), 233–246. 10.1002/ana.24995

Ball, G., Srinivasan, L., Aljabar, P., Counsell, S. J., Durighel, G., Hajnal, J. V., Rutherford, M. A., & Edwards, A. D. (2013). Development of cortical microstructure in the preterm human brain. Proceedings of the National Academy of Sciences, 110(23), 9541–9546. 10.1073/pnas.1301652110

Barbas, H. (2015). General Cortical and Special Prefrontal Connections: Principles from Structure to Function. Annual Review of Neuroscience, 38(1), 269–289. 10.1146/annurev-neuro-071714-033936

Basser, P. J., Mattiello, J., & Lebihan, D. (1994). Estimation of the Effective Self-Diffusion Tensor from the NMR Spin Echo. Journal of Magnetic Resonance, Series B, 103(3), 247–254. 10.1006/jmrb.1994.1037

Batalle, D., Hughes, E. J., Zhang, H., Tournier, J.-D., Tusor, N., Aljabar, P., Wali, L., Alexander, D. C., Hajnal, J. V., Nosarti, C., Edwards, A. D., & Counsell, S. J. (2017). Early development of structural networks and the impact of prematurity on brain connectivity. NeuroImage, 149, 379–392. 10.1016/j.neuroimage.2017.01.065

Batalle, D., O’Muircheartaigh, J., Makropoulos, A., Kelly, C. J., Dimitrova, R., Hughes, E. J., Hajnal, J. V., Zhang, H., Alexander, D. C., Edwards, A. D., & Counsell, S. J. (2019). Different patterns of cortical maturation before and after 38 weeks gestational age demonstrated by diffusion MRI in vivo. NeuroImage, 185, 764–775. 10.1016/j.neuroimage.2018.05.046

Baum, G. L., Cui, Z., Roalf, D. R., Ciric, R., Betzel, R. F., Larsen, B., Cieslak, M., Cook, P. A., Xia, C. H., Moore, T. M., Ruparel, K., Oathes, D. J., Alexander-Bloch, A. F., Shinohara, R. T., Raznahan, A., Gur, R. E., Gur, R. C., Bassett, D. S., & Satterthwaite, T. D. (2020). Development of structure– function coupling in human brain networks during youth. Proceedings of the National Academy of Sciences, 117(1), 771–778. 10.1073/pnas.1912034117

Benavente-Fernández, I., Synnes, A., Grunau, R. E., Chau, V., Ramraj, C., Glass, T., Cayam-Rand, D., Siddiqi, A., & Miller, S. P. (2019). Association of Socioeconomic Status and Brain Injury With Neurodevelopmental Outcomes of Very Preterm Children. JAMA Network Open, 2(5), e192914. 10.1001/jamanetworkopen.2019.2914

Brenner, R. G., Wheelock, M. D., Neil, J. J., & Smyser, C. D. (2021). Structural and functional connectivity in premature neonates. Seminars in Perinatology, 45(7), 151473. 10.1016/j.semperi.2021.151473

Brodmann, K. (1908). Beiträge zur histologischen Lokalisation der Grosshirnrinde. VI. Mitteilung: Die Cortexgliederung des Menschen. Journal Für Psychologie Und Neurologie, 10, 231–246.

Bystron, I., Blakemore, C., & Rakic, P. (2008). Development of the human cerebral cortex: Boulder Committee revisited. Nature Reviews Neuroscience, 9(2), 110–122. 10.1038/nrn2252

Cadwell, C. R., Bhaduri, A., Mostajo-Radji, M. A., Keefe, M. G., & Nowakowski, T. J. (2019). Development and Arealization of the Cerebral Cortex. Neuron, 103(6), 980–1004. 10.1016/j.neuron.2019.07.009

Cao, M., Huang, H., & He, Y. (2017). Developmental Connectomics from Infancy through Early Childhood. Trends in Neurosciences, 40(8), 494–506. 10.1016/j.tins.2017.06.003

Chiarelli, A. M., Sestieri, C., Navarra, R., Wise, R. G., & Caulo, M. (2021). Distinct effects of prematurity on MRI metrics of brain functional connectivity, activity, and structure: Univariate and multivariate analyses. Human Brain Mapping, 42(11), 3593–3607. 10.1002/hbm.25456

Christiaens, D., Cordero-Grande, L., Pietsch, M., Hutter, J., Price, A. N., Hughes, E. J., Vecchiato, K., Deprez, M., Edwards, A. D., Hajnal, J. V., & Tournier, J.-D. (2021). Scattered slice SHARD reconstruction for motion correction in multi-shell diffusion MRI. NeuroImage, 225, 117437. 10.1016/j.neuroimage.2020.117437

Ciarrusta, J., Christiaens, D., Fitzgibbon, S. P., Dimitrova, R., Hutter, J., Hughes, E., Duff, E., Price, A. N., Cordero-Grande, L., Tournier, J.-D., Rueckert, D., Hajnal, J. V., Arichi, T., McAlonan, G., Edwards, A. D., & Batalle, D. (2022). The developing brain structural and functional connectome fingerprint. Developmental Cognitive Neuroscience, 55, 101117. 10.1016/j.dcn.2022.101117

Cordero-Grande, L., Christiaens, D., Hutter, J., Price, A. N., & Hajnal, J. V. (2019). Complex diffusion-weighted image estimation via matrix recovery under general noise models. NeuroImage, 200, 391–404. 10.1016/j.neuroimage.2019.06.039

Dall’Orso, S., Arichi, T., Fitzgibbon, S. P., Edwards, A. D., Burdet, E., & Muceli, S. (2022). Development of functional organization within the sensorimotor network across the perinatal period. Human Brain Mapping, 43(7), 2249–2261. 10.1002/hbm.25785

de Reus, M. A., Saenger, V. M., Kahn, R. S., & van den Heuvel, M. P. (2014). An edge-centric perspective on the human connectome: link communities in the brain. Philosophical Transactions of the Royal Society B: Biological Sciences, 369(1653), 20130527. 10.1098/rstb.2013.0527

Denisova, K. (2019). Age attenuates noise and increases symmetry of head movements during sleep resting-state fMRI in healthy neonates, infants, and toddlers. Infant Behavior and Development, 57, 101317. 10.1016/j.infbeh.2019.03.008

Dimitrova, R., Pietsch, M., Ciarrusta, J., Fitzgibbon, S. P., Williams, L. Z. J., Christiaens, D., Cordero-Grande, L., Batalle, D., Makropoulos, A., Schuh, A., Price, A. N., Hutter, J., Teixeira, R. P., Hughes, E., Chew, A., Falconer, S., Carney, O., Egloff, A., Tournier, J.-D.,…O’Muircheartaigh, J. (2021). Preterm birth alters the development of cortical microstructure and morphology at term-equivalent age. NeuroImage, 243, 118488. 10.1016/j.neuroimage.2021.118488

Doria, V., Beckmann, C. F., Arichi, T., Merchant, N., Groppo, M., Turkheimer, F. E., Counsell, S. J., Murgasova, M., Aljabar, P., Nunes, R. G., Larkman, D. J., Rees, G., & Edwards, A. D. (2010). Emergence of resting state networks in the preterm human brain. Proceedings of the National Academy of Sciences, 107(46), 20015–20020. 10.1073/pnas.1007921107

Dubois, J., Alison, M., Counsell, S. J., Hertz-Pannier, L., Hüppi, P. S., & Benders, M. J. N. L. (2021). MRI of the Neonatal Brain: A Review of Methodological Challenges and Neuroscientific Advances. Journal of Magnetic Resonance Imaging, 53(5), 1318–1343. 10.1002/jmri.27192

Eaton-Rosen, Z., Melbourne, A., Orasanu, E., Cardoso, M. J., Modat, M., Bainbridge, A., Kendall, G. S., Robertson, N. J., Marlow, N., & Ourselin, S. (2015). Longitudinal measurement of the developing grey matter in preterm subjects using multi-modal MRI. NeuroImage, 111, 580–589. 10.1016/j.neuroimage.2015.02.010

Eaton-Rosen, Z., Scherrer, B., Melbourne, A., Ourselin, S., Neil, J. J., & Warfield, S. K. (2017). Investigating the maturation of microstructure and radial orientation in the preterm human cortex with diffusion MRI. NeuroImage, 162, 65–72. 10.1016/j.neuroimage.2017.08.013

Edwards, A. D., Rueckert, D., Smith, S. M., Abo Seada, S., Alansary, A., Almalbis, J., Allsop, J., Andersson, J., Arichi, T., Arulkumaran, S., Bastiani, M., Batalle, D., Baxter, L., Bozek, J., Braithwaite, E., Brandon, J., Carney, O., Chew, A., Christiaens, D.,…Hajnal, J. V. (2022). The Developing Human Connectome Project Neonatal Data Release. Frontiers in Neuroscience, 16. 10.3389/fnins.2022.886772

Eyre, M., Fitzgibbon, S. P., Ciarrusta, J., Cordero-Grande, L., Price, A. N., Poppe, T., Schuh, A., Hughes, E., O’Keeffe, C., Brandon, J., Cromb, D., Vecchiato, K., Andersson, J., Duff, E. P., Counsell, S. J., Smith, S. M., Rueckert, D., Hajnal, J. V, Arichi, T.,…Edwards, A. D. (2021). The Developing Human Connectome Project: typical and disrupted perinatal functional connectivity. Brain, 144(7), 2199–2213. 10.1093/brain/awab118

Fan, Y., Shi, F., Smith, J. K., Lin, W., Gilmore, J. H., & Shen, D. (2011). Brain anatomical networks in early human brain development. NeuroImage, 54(3), 1862–1871. 10.1016/j.neuroimage.2010.07.025

Fenchel, D., Dimitrova, R., Seidlitz, J., Robinson, E. C., Batalle, D., Hutter, J., Christiaens, D., Pietsch, M., Brandon, J., Hughes, E. J., Allsop, J., O’Keeffe, C., Price, A. N., Cordero-Grande, L., Schuh, A., Makropoulos, A., Passerat-Palmbach, J., Bozek, J., Rueckert, D.,…O’Muircheartaigh, J. (2020). Fench et Development of Microstructural and Morphological Cortical Profiles in the Neonatal Brain. Cerebral Cortex, 30(11), 5767–5779. 10.1093/cercor/bhaa150

Fitzgibbon, S. P., Harrison, S. J., Jenkinson, M., Baxter, L., Robinson, E. C., Bastiani, M., Bozek, J., Karolis, V., Cordero Grande, L., Price, A. N., Hughes, E., Makropoulos, A., Passerat-Palmbach, J., Schuh, A., Gao, J., Farahibozorg, S. R., O’Muircheartaigh, J., Ciarrusta, J., O’Keeffe, C.,…Andersson, J. (2020). The developing Human Connectome Project (dHCP) automated resting-state functional processing framework for newborn infants. NeuroImage, 223(August), 117303. 10.1016/j.neuroimage.2020.117303

Fransson, P., Skiöld, B., Engström, M., Hallberg, B., Mosskin, M., Åden, U., Lagercrantz, H., & Blennow, M. (2009). Spontaneous Brain Activity in the Newborn Brain During Natural Sleep— An fMRI Study in Infants Born at Full Term. Pediatric Research, 66(3), 301–305. 10.1203/PDR.0b013e3181b1bd84

Friedrichs-Maeder, C. L., Griffa, A., Schneider, J., Hüppi, P. S., Truttmann, A., & Hagmann, P. (2017). Exploring the role of white matter connectivity in cortex maturation. PLOS ONE, 12(5), e0177466. 10.1371/journal.pone.0177466

Fukutomi, H., Glasser, M. F., Zhang, H., Autio, J. A., Coalson, T. S., Okada, T., Togashi, K., Van Essen, D. C., & Hayashi, T. (2018). Neurite imaging reveals microstructural variations in human cerebral cortical gray matter. NeuroImage, 182, 488–499. 10.1016/j.neuroimage.2018.02.017

Galdi, P., Blesa, M., Stoye, D. Q., Sullivan, G., Lamb, G. J., Quigley, A. J., Thrippleton, M. J., Bastin, M. E., & Boardman, J. P. (2020). Neonatal morphometric similarity mapping for predicting brain age and characterizing neuroanatomic variation associated with preterm birth. NeuroImage: Clinical, 25, 102195. 10.1016/j.nicl.2020.102195

Gao, W., Alcauter, S., Elton, A., Hernandez-Castillo, C. R., Smith, J. K., Ramirez, J., & Lin, W. (2015). Functional Network Development During the First Year: Relative Sequence and Socioeconomic Correlations. Cerebral Cortex, 25(9), 2919–2928. 10.1093/cercor/bhu088

Geng, X., Li, G., Lu, Z., Gao, W., Wang, L., Shen, D., Zhu, H., & Gilmore, J. H. (2016). Structural and Maturational Covariance in Early Childhood Brain Development. *Cerebral Cortex*, bhw022. 10.1093/cercor/bhw022

Gilmore, J. H., Knickmeyer, R. C., & Gao, W. (2018). Imaging structural and functional brain development in early childhood. Nature Reviews Neuroscience, 19(3), 123–137. 10.1038/nrn.2018.1

Gilmore, J. H., Shi, F., Woolson, S. L., Knickmeyer, R. C., Short, S. J., Lin, W., Zhu, H., Hamer, R. M., Styner, M., & Shen, D. (2012). Longitudinal Development of Cortical and Subcortical Gray Matter from Birth to 2 Years. Cerebral Cortex, 22(11), 2478–2485. 10.1093/cercor/bhr327

Gondová, A., Neumane, S., Leprince, Y., Mangin, J.-F., Arichi, T., & Dubois, J. (2023). Predicting neurodevelopmental outcomes from neonatal cortical microstructure: A conceptual replication study. Neuroimage: Reports, 3(2), 100170. 10.1016/j.ynirp.2023.100170

Goulas, A., Uylings, H. B. M., & Hilgetag, C. C. (2017). Principles of ipsilateral and contralateral cortico-cortical connectivity in the mouse. Brain Structure and Function, 222(3), 1281–1295. 10.1007/s00429-016-1277-y

Goulas, A., Werner, R., Beul, S. F., Saering, D., Heuvel, M. P., Triarhou, L. C., & Hilgetag, C. C. (2016). Cytoarchitectonic similarity is a wiring principle of the human connectome. BioRxiv. 10.1101/068254

Grayson, D. S., Ray, S., Carpenter, S., Iyer, S., Dias, T. G. C., Stevens, C., Nigg, J. T., & Fair, D. A. (2014). Structural and Functional Rich Club Organization of the Brain in Children and Adults. PLoS ONE, 9(2), e88297. 10.1371/journal.pone.0088297

Guerrero, J. M., Adluru, N., Bendlin, B. B., Goldsmith, H. H., Schaefer, S. M., Davidson, R. J., Kecskemeti, S. R., Zhang, H., & Alexander, A. L. (2019). Optimizing the intrinsic parallel diffusivity in NODDI: An extensive empirical evaluation. PLOS ONE, 14(9), e0217118. 10.1371/journal.pone.0217118

Hagmann, P., Sporns, O., Madan, N., Cammoun, L., Pienaar, R., Wedeen, V. J., Meuli, R., Thiran, J.-P., & Grant, P. E. (2010). White matter maturation reshapes structural connectivity in the late developing human brain. Proceedings of the National Academy of Sciences, 107(44), 19067– 19072. 10.1073/pnas.1009073107

Hernandez-Fernandez, M., Reguly, I., Jbabdi, S., Giles, M., Smith, S., & Sotiropoulos, S. N. (2019). Using GPUs to accelerate computational diffusion MRI: From microstructure estimation to tractography and connectomes. NeuroImage, 188, 598–615. 10.1016/j.neuroimage.2018.12.015

Hoff, G. E. A.-J., Van den Heuvel, M. P., Benders, M. J. N. L., Kersbergen, K. J., & De Vries, L. S. (2013). On development of functional brain connectivity in the young brain. Frontiers in Human Neuroscience, 7. 10.3389/fnhum.2013.00650

Hughes, E. J., Winchman, T., Padormo, F., Teixeira, R., Wurie, J., Sharma, M., Fox, M., Hutter, J., Cordero-Grande, L., Price, A. N., Allsop, J., Bueno-Conde, J., Tusor, N., Arichi, T., Edwards, A. D., Rutherford, M. A., Counsell, S. J., & Hajnal, J. V. (2017). A dedicated neonatal brain imaging system. Magnetic Resonance in Medicine, 78(2), 794–804. 10.1002/mrm.26462

Hutter, J., Tournier, J. D., Price, A. N., Cordero-Grande, L., Hughes, E. J., Malik, S., Steinweg, J., Bastiani, M., Sotiropoulos, S. N., Jbabdi, S., Andersson, J., Edwards, A. D., & Hajnal, J. V. (2018). Time-efficient and flexible design of optimized multishell HARDI diffusion. Magnetic Resonance in Medicine, 79(3), 1276–1292. 10.1002/mrm.26765

Jakab, A., Schwartz, E., Kasprian, G., Gruber, G. M., Prayer, D., SchÃ¶pf, V., & Langs, G. (2014). Fetal functional imaging portrays heterogeneous development of emerging human brain networks. Frontiers in Human Neuroscience, 8. 10.3389/fnhum.2014.00852

Jelescu, I. O., Veraart, J., Adisetiyo, V., Milla, S. S., Novikov, D. S., & Fieremans, E. (2015). One diffusion acquisition and different white matter models: How does microstructure change in human early development based on WMTI and NODDI? NeuroImage, 107, 242–256. 10.1016/j.neuroimage.2014.12.009

Keunen, K., Counsell, S. J., & Benders, M. J. N. L. (2017). The emergence of functional architecture during early brain development. NeuroImage, 160, 2–14. 10.1016/j.neuroimage.2017.01.047

Khundrakpam, B. S., Lewis, J. D., Zhao, L., Chouinard-Decorte, F., & Evans, A. C. (2016). Brain connectivity in normally developing children and adolescents. NeuroImage, 134, 192–203. 10.1016/j.neuroimage.2016.03.062

Khundrakpam, B. S., Reid, A., Brauer, J., Carbonell, F., Lewis, J., Ameis, S., Karama, S., Lee, J., Chen, Z., Das, S., Evans, A. C., Ball, W. S., Byars, A. W., Schapiro, M., Bommer, W., Carr, A., German, A., Dunn, S., Rivkin, M. J.,…O’Neill, J. (2013). Developmental Changes in Organization of Structural Brain Networks. Cerebral Cortex, 23(9), 2072–2085. 10.1093/cercor/bhs187

King, D. J., & Wood, A. G. (2020). Clinically feasible brain morphometric similarity network construction approaches with restricted magnetic resonance imaging acquisitions. Network Neuroscience, 4(1), 274–291. 10.1162/netn_a_00123

Klein, A., & Tourville, J. (2012). 101 Labeled Brain Images and a Consistent Human Cortical Labeling Protocol. Frontiers in Neuroscience, 6. 10.3389/fnins.2012.00171

Kostović, I., Radoš, M., Kostović-Srzentić, M., & Krsnik, Ž. (2021). Fundamentals of the Development of Connectivity in the Human Fetal Brain in Late Gestation: From 24 Weeks Gestational Age to Term. Journal of Neuropathology & Experimental Neurology, 80(5), 393–414. 10.1093/jnen/nlab024

Kostović, I., Sedmak, G., & Judaš, M. (2019). Neural histology and neurogenesis of the human fetal and infant brain. NeuroImage, 188, 743–773. 10.1016/j.neuroimage.2018.12.043

Kulikova, S., Hertz-Pannier, L., Dehaene-Lambertz, G., Buzmakov, A., Poupon, C., & Dubois, J. (2015). Multi-parametric evaluation of the white matter maturation. Brain Structure and Function, 220(6), 3657–3672. 10.1007/s00429-014-0881-y

Larivière, S., Vos de Wael, R., Hong, S.-J., Paquola, C., Tavakol, S., Lowe, A. J., Schrader, D. V, & Bernhardt, B. C. (2020). Multiscale Structure–Function Gradients in the Neonatal Connectome. Cerebral Cortex, 30(1), 47–58. 10.1093/cercor/bhz069

Lebenberg, J., Mangin, J.-F., Thirion, B., Poupon, C., Hertz-Pannier, L., Leroy, F., Adibpour, P., Dehaene-Lambertz, G., & Dubois, J. (2019). Mapping the asynchrony of cortical maturation in the infant brain: A MRI multi-parametric clustering approach. NeuroImage, 185, 641–653. 10.1016/j.neuroimage.2018.07.022

Lyall, A. E., Shi, F., Geng, X., Woolson, S., Li, G., Wang, L., Hamer, R. M., Shen, D., & Gilmore, J. H. (2015). Dynamic Development of Regional Cortical Thickness and Surface Area in Early Childhood. Cerebral Cortex, 25(8), 2204–2212. 10.1093/cercor/bhu027

Makropoulos, A., Gousias, I. S., Ledig, C., Aljabar, P., Serag, A., Hajnal, J. V., Edwards, A. D., Counsell, S. J., & Rueckert, D. (2014). Automatic Whole Brain MRI Segmentation of the Developing Neonatal Brain. IEEE Transactions on Medical Imaging, 33(9), 1818–1831. 10.1109/TMI.2014.2322280

Makropoulos, A., Robinson, E. C., Schuh, A., Wright, R., Fitzgibbon, S., Bozek, J., Counsell, S. J., Steinweg, J., Vecchiato, K., Passerat-Palmbach, J., Lenz, G., Mortari, F., Tenev, T., Duff, E. P., Bastiani, M., Cordero-Grande, L., Hughes, E., Tusor, N., Tournier, J. D.,…Rueckert, D. (2018). The developing human connectome project: A minimal processing pipeline for neonatal cortical surface reconstruction. NeuroImage, 173(April 2017), 88–112. 10.1016/j.neuroimage.2018.01.054

Maugeri, L., Moraschi, M., Summers, P., Favilla, S., Mascali, D., Cedola, A., Porro, C. A., Giove, F., & Fratini, M. (2018). Assessing denoising strategies to increase signal to noise ratio in spinal cord and in brain cortical and subcortical regions. Journal of Instrumentation, 13(02), C02028– C02028. 10.1088/1748-0221/13/02/C02028

Monson, B. B., Eaton-Rosen, Z., Kapur, K., Liebenthal, E., Brownell, A., Smyser, C. D., Rogers, C. E., Inder, T. E., Warfield, S. K., & Neil, J. J. (2018). Differential Rates of Perinatal Maturation of Human Primary and Nonprimary Auditory Cortex. Eneuro, 5(1), ENEURO.0380-17.2017. 10.1523/ENEURO.0380-17.2017

Moreno-Dominguez, D., Anwander, A., & Knösche, T. R. (2014). A hierarchical method for whole-brain connectivity-based parcellation. Human Brain Mapping, 35(10), 5000–5025. 10.1002/hbm.22528

Mukherjee, P., Miller, J. H., Shimony, J. S., Conturo, T. E., Lee, B. C. P., Almli, C. R., & McKinstry, R. C. (2001). Normal Brain Maturation during Childhood: Developmental Trends Characterized with Diffusion-Tensor MR Imaging. Radiology, 221(2), 349–358. 10.1148/radiol.2212001702

Neil, J. J., & Smyser, C. D. (2018). Recent advances in the use of MRI to assess early human cortical development. Journal of Magnetic Resonance, 293, 56–69. 10.1016/j.jmr.2018.05.013

Neumane, S., Gondova, A., Leprince, Y., Hertz-Pannier, L., Arichi, T., & Dubois, J. (2022). Early structural connectivity within the sensorimotor network: Deviations related to prematurity and association to neurodevelopmental outcome. Frontiers in Neuroscience, 16. 10.3389/fnins.2022.932386

Nie, J., Li, G., Wang, L., Shi, F., Lin, W., Gilmore, J. H., & Shen, D. (2014). Longitudinal development of cortical thickness, folding, and fiber density networks in the first 2 years of life. Human Brain Mapping, 35(8), 3726–3737. 10.1002/hbm.22432

Ouyang, M., Dubois, J., Yu, Q., Mukherjee, P., & Huang, H. (2019). Delineation of early brain development from fetuses to infants with diffusion MRI and beyond. NeuroImage, 185, 836– 850. 10.1016/j.neuroimage.2018.04.017

Ouyang, M., Jeon, T., Sotiras, A., Peng, Q., Mishra, V., Halovanic, C., Chen, M., Chalak, L., Rollins, N., Roberts, T. P. L., Davatzikos, C., & Huang, H. (2019). Differential cortical microstructural maturation in the preterm human brain with diffusion kurtosis and tensor imaging. Proceedings of the National Academy of Sciences, 116(10), 4681–4688. 10.1073/pnas.1812156116

Price, A. N. (2015). Accelerated Neonatal fMRI Using Multiband EPI.

Risk, B. B., Murden, R. J., Wu, J., Nebel, M. B., Venkataraman, A., Zhang, Z., & Qiu, D. (2021). Which multiband factor should you choose for your resting-state fMRI study? NeuroImage, 234, 117965. 10.1016/j.neuroimage.2021.117965

Romero-Garcia, R., Whitaker, K. J., Váša, F., Seidlitz, J., Shinn, M., Fonagy, P., Dolan, R. J., Jones, P. B., Goodyer, I. M., Bullmore, E. T., & Vértes, P. E. (2018). Structural covariance networks are coupled to expression of genes enriched in supragranular layers of the human cortex. NeuroImage, 171, 256–267. 10.1016/j.neuroimage.2017.12.060

Scheinost, D., Kwon, S. H., Shen, X., Lacadie, C., Schneider, K. C., Dai, F., Ment, L. R., & Constable, R. T. (2016). Preterm birth alters neonatal, functional rich club organization. Brain Structure and Function, 221(6), 3211–3222. 10.1007/s00429-015-1096-6

Seidlitz, J., Váša, F., Shinn, M., Romero-Garcia, R., Whitaker, K. J., Vértes, P. E., Wagstyl, K., Kirkpatrick Reardon, P., Clasen, L., Liu, S., Messinger, A., Leopold, D. A., Fonagy, P., Dolan, R. J., Jones, P. B., Goodyer, I. M., Raznahan, A., & Bullmore, E. T. (2018). Morphometric Similarity Networks Detect Microscale Cortical Organization and Predict Inter-Individual Cognitive Variation. Neuron, 97(1), 231–247.e7. 10.1016/j.neuron.2017.11.039

Smith, S. M., & Beckmann, C. F. (2017). Introduction to Resting State fMRI Functional Connectivity.

Smyser, C. D., Inder, T. E., Shimony, J. S., Hill, J. E., Degnan, A. J., Snyder, A. Z., & Neil, J. J. (2010). Longitudinal Analysis of Neural Network Development in Preterm Infants. Cerebral Cortex, 20(12), 2852–2862. 10.1093/cercor/bhq035

Smyser, T. A., Smyser, C. D., Rogers, C. E., Gillespie, S. K., Inder, T. E., & Neil, J. J. (2016). Cortical Gray and Adjacent White Matter Demonstrate Synchronous Maturation in Very Preterm Infants. Cerebral Cortex, 26(8), 3370–3378. 10.1093/cercor/bhv164

Takahashi, E., Folkerth, R. D., Galaburda, A. M., & Grant, P. E. (2012). Emerging Cerebral Connectivity in the Human Fetal Brain: An MR Tractography Study. Cerebral Cortex, 22(2), 455–464. 10.1093/cercor/bhr126

Taymourtash, A., Schwartz, E., Nenning, K.-H., Sobotka, D., Licandro, R., Glatter, S., Diogo, M. C., Golland, P., Grant, E., Prayer, D., Kasprian, G., & Langs, G. (2023). Fetal development of functional thalamocortical and cortico–cortical connectivity. Cerebral Cortex, 33(9), 5613–5624. 10.1093/cercor/bhac446

Thirion, B., Varoquaux, G., Dohmatob, E., & Poline, J.-B. (2014). Which fMRI clustering gives good brain parcellations? Frontiers in Neuroscience, 8. 10.3389/fnins.2014.00167

Thomason, M. E., Grove, L. E., Lozon, T. A., Vila, A. M., Ye, Y., Nye, M. J., Manning, J. H., Pappas, A., Hernandez-Andrade, E., Yeo, L., Mody, S., Berman, S., Hassan, S. S., & Romero, R. (2015). Age-related increases in long-range connectivity in fetal functional neural connectivity networks in utero. Developmental Cognitive Neuroscience, 11, 96–104. 10.1016/j.dcn.2014.09.001

Toulmin, H., Beckmann, C. F., O’Muircheartaigh, J., Ball, G., Nongena, P., Makropoulos, A., Ederies, A., Counsell, S. J., Kennea, N., Arichi, T., Tusor, N., Rutherford, M. A., Azzopardi, D., Gonzalez-Cinca, N., Hajnal, J. V., & Edwards, A. D. (2015). Specialization and integration of functional thalamocortical connectivity in the human infant. Proceedings of the National Academy of Sciences, 112(20), 6485–6490. 10.1073/pnas.1422638112

Turk, E., van den Heuvel, M. I., Benders, M. J., de Heus, R., Franx, A., Manning, J. H., Hect, J. L., Hernandez-Andrade, E., Hassan, S. S., Romero, R., Kahn, R. S., Thomason, M. E., & van den Heuvel, M. P. (2019). Functional Connectome of the Fetal Brain. The Journal of Neuroscience, 39(49), 9716–9724. 10.1523/JNEUROSCI.2891-18.2019

van den Heuvel, M. P., & Hulshoff Pol, H. E. (2010). Exploring the brain network: A review on resting-state fMRI functional connectivity. European Neuropsychopharmacology, 20(8), 519–534. 10.1016/j.euroneuro.2010.03.008

van den Heuvel, M. P., Kersbergen, K. J., de Reus, M. A., Keunen, K., Kahn, R. S., Groenendaal, F., de Vries, L. S., & Benders, M. J. N. L. (2015). The Neonatal Connectome During Preterm Brain Development. Cerebral Cortex, 25(9), 3000–3013. 10.1093/cercor/bhu095

Vanes, L., Fenn-Moltu, S., Hadaya, L., Fitzgibbon, S., Cordero-Grande, L., Price, A., Chew, A., Falconer, S., Arichi, T., Counsell, S. J., Hajnal, J. V., Batalle, D., Edwards, A. D., & Nosarti, C. (2023). Longitudinal neonatal brain development and socio-demographic correlates of infant outcomes following preterm birth. Developmental Cognitive Neuroscience, 61, 101250. 10.1016/j.dcn.2023.101250

Váša, F., Seidlitz, J., Romero-Garcia, R., Whitaker, K. J., Rosenthal, G., Vértes, P. E., Shinn, M., Alexander-Bloch, A., Fonagy, P., Dolan, R. J., Jones, P. B., Goodyer, I. M., Sporns, O., & Bullmore, E. T. (2018). Adolescent Tuning of Association Cortex in Human Structural Brain Networks. Cerebral Cortex, 28(1), 281–294. 10.1093/cercor/bhx249

Vasung, L., Raguz, M., Kostovic, I., & Takahashi, E. (2017). Spatiotemporal Relationship of Brain Pathways during Human Fetal Development Using High-Angular Resolution Diffusion MR Imaging and Histology. Frontiers in Neuroscience, 11. 10.3389/fnins.2017.00348

von Economo, C. F., & Koskinas, G. N. (1925). Die cytoarchitektonik der hirnrinde des erwachsenen menschen. J. Springer.

Williams, L. Z. J., Fitzgibbon, S. P., Bozek, J., Winkler, A. M., Dimitrova, R., Poppe, T., Schuh, A., Makropoulos, A., Cupitt, J., O’Muircheartaigh, J., Duff, E. P., Cordero-Grande, L., Price, A. N., Hajnal, J. V., Rueckert, D., Smith, S. M., Edwards, A. D., & Robinson, E. C. (2023). Structural and functional asymmetry of the neonatal cerebral cortex. Nature Human Behaviour, 7(6), 942–955. 10.1038/s41562-023-01542-8

Wilson, S., Pietsch, M., Cordero-Grande, L., Christiaens, D., Uus, A., Karolis, V. R., Kyriakopoulou, V., Colford, K., Price, A. N., Hutter, J., Rutherford, M. A., Hughes, E. J., Counsell, S. J., Tournier, J.-D., Hajnal, J. V, Edwards, A. D., O’Muircheartaigh, J., & Arichi, T. (2023). Spatiotemporal tissue maturation of thalamocortical pathways in the human fetal brain. ELife, 12. 10.7554/eLife.83727

Wilson, S., Pietsch, M., Cordero-Grande, L., Price, A. N., Hutter, J., Xiao, J., McCabe, L., Rutherford, M. A., Hughes, E. J., Counsell, S. J., Tournier, J.-D., Arichi, T., Hajnal, J. V., Edwards, A. D., Christiaens, D., & O’Muircheartaigh, J. (2021). Development of human white matter pathways in utero over the second and third trimester. Proceedings of the National Academy of Sciences, 118(20). 10.1073/pnas.2023598118

Yee, Y., Fernandes, D. J., French, L., Ellegood, J., Cahill, L. S., Vousden, D. A., Spencer Noakes, L., Scholz, J., van Eede, M. C., Nieman, B. J., Sled, J. G., & Lerch, J. P. (2018). Structural covariance of brain region volumes is associated with both structural connectivity and transcriptomic similarity. NeuroImage, 179, 357–372. 10.1016/j.neuroimage.2018.05.028

Yeo, B. T., Krienen, F. M., Sepulcre, J., Sabuncu, M. R., Lashkari, D., Hollinshead, M., Roffman, J. L., Smoller, J. W., Zöllei, L., Polimeni, J. R., Fischl, B., Liu, H., & Buckner, R. L. (2011). The organization of the human cerebral cortex estimated by intrinsic functional connectivity. Journal of Neurophysiology, 106(3), 1125–1165. 10.1152/jn.00338.2011

Yu, Q., Ouyang, A., Chalak, L., Jeon, T., Chia, J., Mishra, V., Sivarajan, M., Jackson, G., Rollins, N., Liu, S., & Huang, H. (2016). Structural Development of Human Fetal and Preterm Brain Cortical Plate Based on Population-Averaged Templates. Cerebral Cortex, 26(11), 4381–4391. 10.1093/cercor/bhv201

Zhang, H., Schneider, T., Wheeler-Kingshott, C. A., & Alexander, D. C. (2012). NODDI: Practical in vivo neurite orientation dispersion and density imaging of the human brain. NeuroImage, 61(4), 1000–1016. 10.1016/j.neuroimage.2012.03.072

Zhang, L., Wang, L., & Zhu, D. (2022). Predicting brain structural network using functional connectivity. Medical Image Analysis, 79, 102463. 10.1016/j.media.2022.102463

