## Supplementary Materials for "Early development and co-evolution of microstructural and functional brain connectomes: A multi-modal MRI study in preterm and full-term infants"

#### *SI1. Cohort description*

**a,**

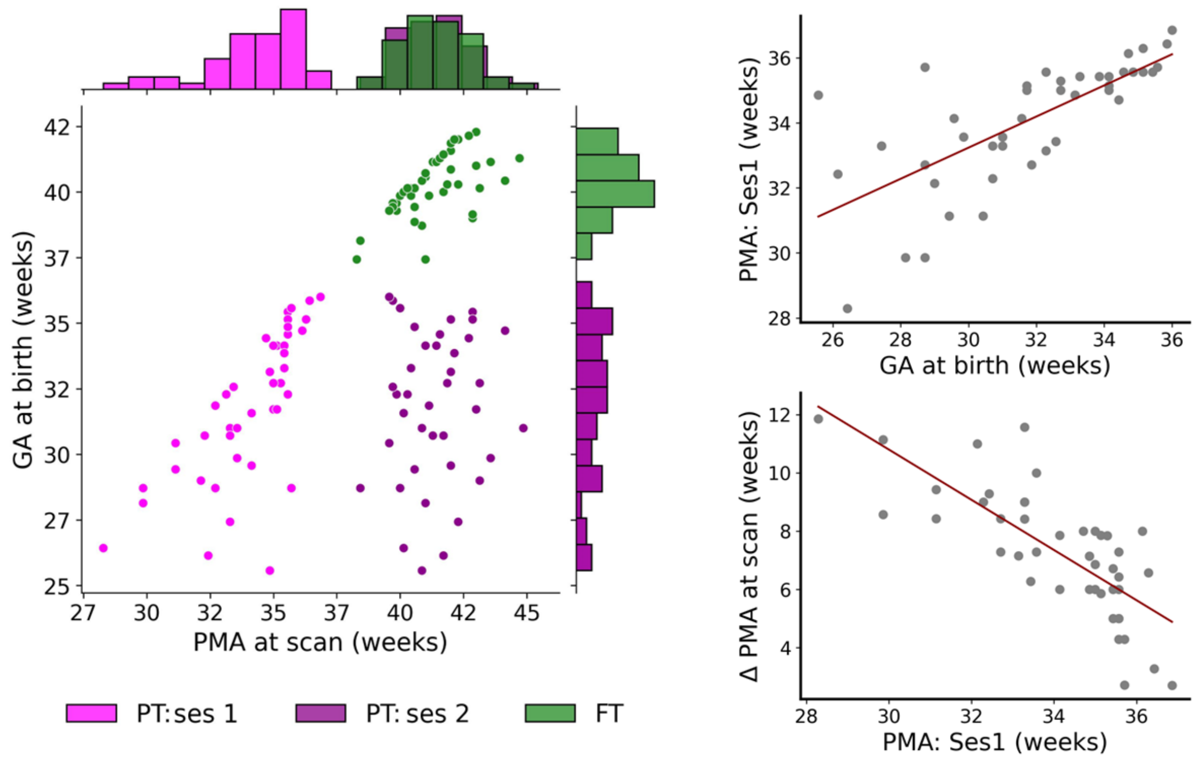


**b,**

| | *FT (N = 45)* | | *PT:ses 1 (N = 45)* | |
|---|---|---|---|---|
| | **Mean (std)** | **Median [range]** | **Mean (std)** | **Median [range]** |
| **GA at birth** | 40.3 (1.17) | 40.1, [37.4,42.3] | 31.8 (2.84) | 32.3, [25.6,36.0] |
| **PMA scan: ses1** | - | - | 34.1 (1.88) | 34.9, [28.3,36.9] |
| **PMA at scan: ses2 (TEA)** | 41.4 (1.41) | 41.3, [38.3,44.7] | 41.4 (1.42) | 41.3, [38.4,44.9] |
| **N (%) male** | 26 (58%) | | 26 (58%) | |

**Radiology score:**

| | *FT* | *PT:ses 1* | *PT:ses2* | |
|---|---|---|---|---|
| **1** | 29 (64%) | 27 (60%) | 19 (42%) | |
| **2** | 11 (24%) | 10 (22%) | 13 (29%) | p=0.17 |
| **3** | 5 (11%) | 8 (18%) | 13 (29%) | |

**Supp. Figure 1.1. a,** Joined distribution of PMA at scan and GA at birth for the 3 infant groups. Right: significant relationship between the GA at birth with PMA at ses1, and PMA: ses1 with delay between PMA at session 1 and 2 (linear regression, $p<0.001$). *Legend:* FT: Full-term group, PT: ses1: preterm group scanned at 1st session, PT: ses2: preterm group scanned at term-equivalent age, GA: gestational age, PMA: post-menstrual age, Δ: difference. **b,** Table summarizing cohort characteristics. *Legend:* FT: Full-term group, PT:ses1: preterm group scanned at 1st session, PT:ses2: preterm group scanned at term-equivalent age, GA: gestational age, PMA: post-menstrual age, N: number, std: standard deviation. dHCP radiological scores: 1: *normal appearance for age*; 2: *incidental findings with unlikely* *significance for clinical outcome or analysis;* 3: *incidental findings with unlikely clinical but possible analysis significance*.

#### *SI2. Data quality*

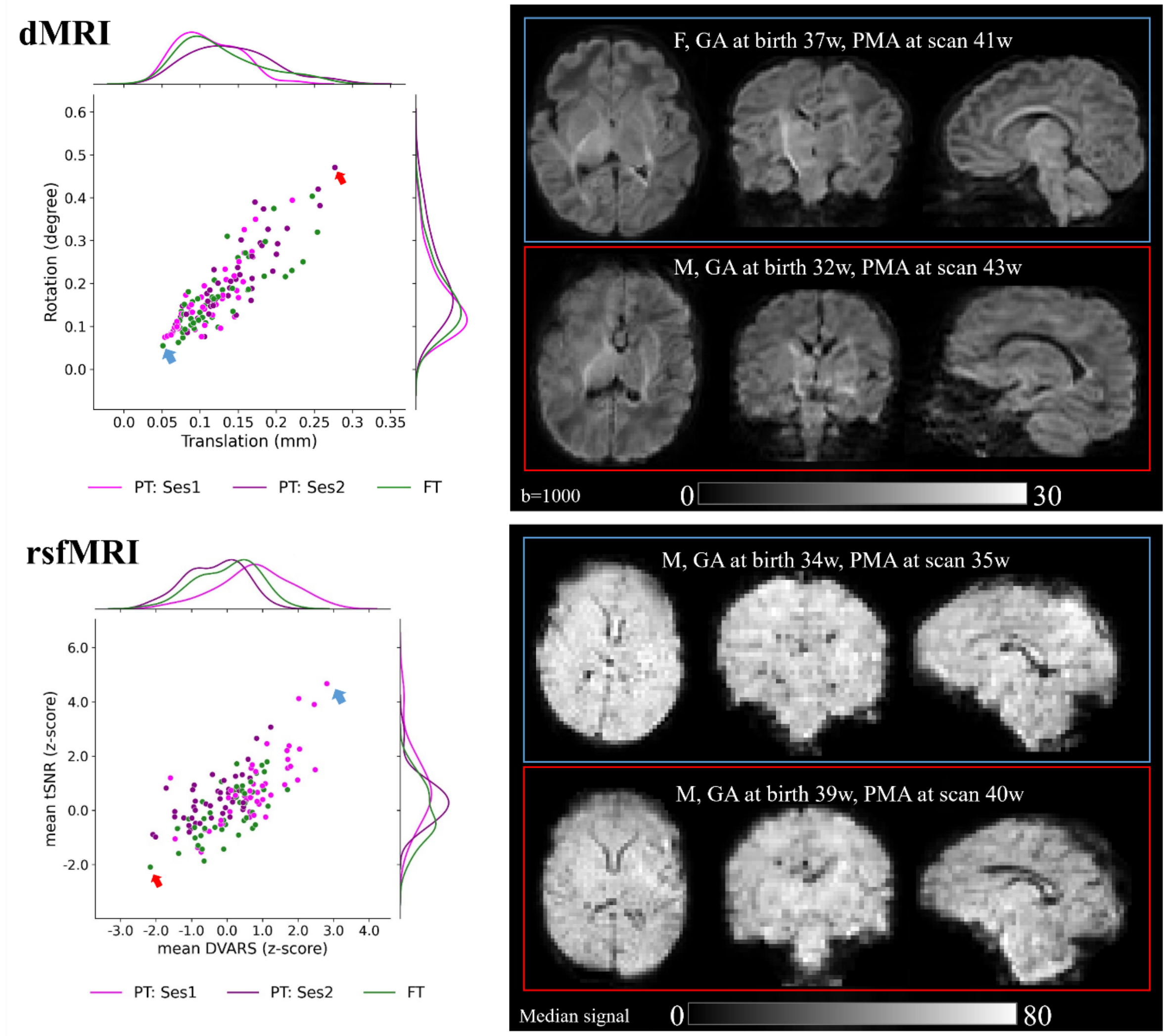


**Supp. Figure 2.1.** Quality control of diffusion and functional data across the cohort (**right**). Example diffusion and resting state functional data for subjects with the highest (in **blue**) and the lowest (in **red**) data quality (**right**). *Legend:* M: male, F: female, tSNR: temporal signal-to-noise ratio, DVARS: the spatial standard deviation of successive difference images. See *Supp. Figure 1* for the remaining abbreviations.

**Supp. Table 2.1.** Quality control metrics of diffusion and functional data. P-values result from paired t-tests. **Legend:** SNR: mean signal-to-noise ratio, tSNR: mean temporal signal-to-noise ratio (converted to Z-score), DVARS: mean spatial standard deviation of successive difference images (converted to Z-score).

| | **PT:ses1** *Z-score, mean (SD)* | **PT:ses2** *Z-score, mean (SD)* | **FT** *Z-score, mean (SD)* | **PT:ses1 vs ses2** *p-values* | **PT:ses1 vs FT** *p-values* | **PT:ses2 vs FT** *p-values* |
|---|---|---|---|---|---|---|
| **Data quality – rs-fMRI** | | | | | | |
| **DVARS** | 0.77 (0.989) | -0.35 (0.778) | 0.01 (0.827) | <0.001 | 0.002 | 0.042 |
| **tSNR** | 0.83 (1.325) | 0.39 (0.833) | -0.06 (0.979) | 0.083 | 0.004 | 0.015 |
| **Data quality - dMRI** | | | | | | |

| | | | | | | |
|---|---|---|---|---|---|---|
| **translation** | 0.11 (0.038) | 0.14 (0.052) | 0.13 (0.051) | 0.005 | 0.187 | 0.188 |
| **rotation** | 0.16 (0.077) | 0.21 (0.096) | 0.17 (0.081) | 0.017 | 0.625 | 0.083 |
| **SNR** | 42.55 (3.449) | 41.37 (2.405) | 41.66 (5.062) | 0.077 | 0.567 | 0.898 |

**Supp. Table 2.2. a,** Group differences for selected variables. Reported p-values result from paired t-tests for continuous variables and $\chi^2$ test for sex, and Wilcoxon Signed Rank Test for the radiology score and pregnancy size. **b,** ANOVA model studying effects of selected variables on the six global median diffusion metrics and mean FC. p-value ≤0.001 [***], ≤0.01 [**], < 0.05 [*]. Legend: M: male, F: female, IMD: index of multiple deprivation.

**a,**

| | **PT group** *mean (SD) [range] / N (%)* | **FT group** *mean (SD) [range] / N (%)* | **PT vs FT** *p-values* |
|---|---|---|---|
| **GA at birth *(weeks)*** | 31.80 (2.838) [25.57; 36.00] | 40.25 (1.171) [37.43; 42.29] | 0.001 |
| **PMA at scan *(weeks)*** | *Ses1*: 34.10 (1.881) [28.29; 36.86]<br>*Ses2*: 41.36 (1.416) [38.43; 44.86] | 41.36 (1.409) [38.29; 44.71]<br>- | 0.001<br>0.625 |
| **Sex** | F: 19 (42%) M: 26 (58%) | F: 19 (42%) M: 26 (58%) | - |
| **Radiology score** | 1: 27 (60%) 2:10 (22%) 3: 8 (18%) | 1: 29 (64%) 2: 11 (24%) 3: 5 (12%) | 0.585 |
| **Mother age *(years)*** | 34.56 (5.044) [24.00; 52.00] | 34.44 (4.731) [24.00; 44.00] | 1.000 |
| **Pregnancy size** | 1: 30 (66%) 2: 15 (34%) | 1: 44 (98%) 2: 1 (2%) | 0.001 |
| **IMD *score*** | 21.12 (11.753) [2.68; 48.30] | 23.95 (10.529) [4.17; 46.37] | 0.956 |

**b,**

| | **FA** | **AD** | **RD** | **MD** | **OD** | **NDI** | **Mean FC** |
|---|---|---|---|---|---|---|---|
| *Group* | *** | *** | *** | *** | *** | *** | *** |
| *Sex* | | | * | * | | * | |
| *Radiology score* | *** | | ** | * | ** | * | |
| *IMD* | ** | * | | | | | |
| *Mother age* | | * | * | * | * | * | |
| *Pregnancy size* | * | | | | | | |
| *Group:IMD* | ** | | | | | | |

**a,**

| | | | |
|---|---|---|---|
| ***Cortical (left/right)*** | ***Frontal*** | Superior frontal<br>Rostral middle frontal<br>Caudal middle frontal<br>Pars opercularis<br>Pars triangularis | Pars orbitalis<br>Lateral orbitofrontal<br>Medial orbitofrontal<br>Precentral<br>Paracentral |
| | ***Parietal*** | Superior parietal<br>Inferior parietal<br>Supramarginal | Postcentral<br>Precuneus |
| | ***Temporal*** | Superior temporal<br>Middle temporal<br>Inferior temporal<br>Fusiform | Transverse temporal<br>Entorhinal<br>Parahippocampal |
| | ***Occipital*** | Lateral occipital<br>Lingual | Cuneus<br>Pericalcarine |
| | ***Cingulate*** | Rostral anterior cingulate<br>Posterior cingulate | Isthmus cingulate<br>Caudal anterior cingulate |
| | ***Insular*** | Insula | |
| ***Subcortical (left/right):*** | | Hippocampus (*hippo*)<br>Amygdala (*amyg*)<br>Thalamus (*thal*)<br>Caudate nucleus (*caud*)<br>Lenticular nucleus (*lenti*)<br>Cerebellum (*cereb*) | |
| ***Subcortical (medial):*** | | Brainstem (*bstem*) | |

**b,**

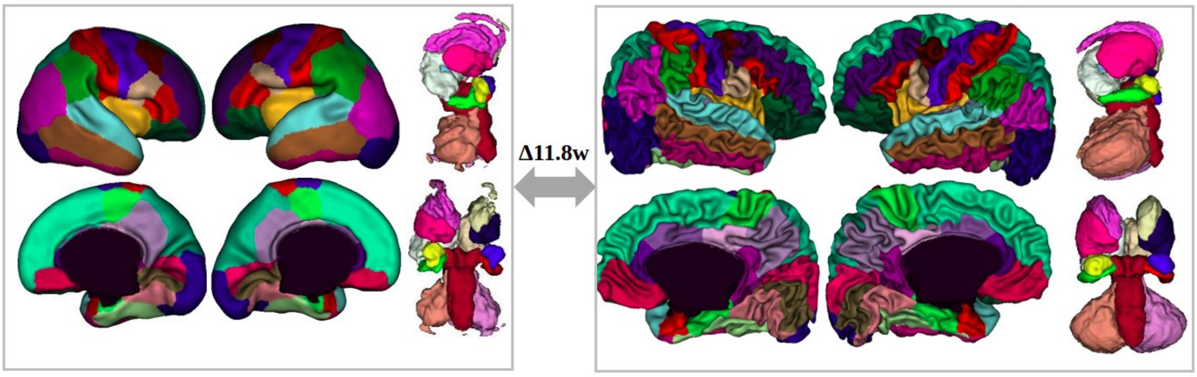


**Supp. Figure 2.2. a,** List of regions of interest (ROIs) used in the study with assignment to the lobes. Color-coding is re-used in the circular plots. We highlight cortical regions considered as primary sensorimotor (SM) system in red, regions of the primary visual (VIS) system in blue. **b,** Example of segmentation for the longitudinal subject with the longest scan delay. **Left**: ses1 scan at 28.3 PMA; **Right**: ses2 acquisition at 40.1 PMA. *Legend:* Δ: difference in PMA.

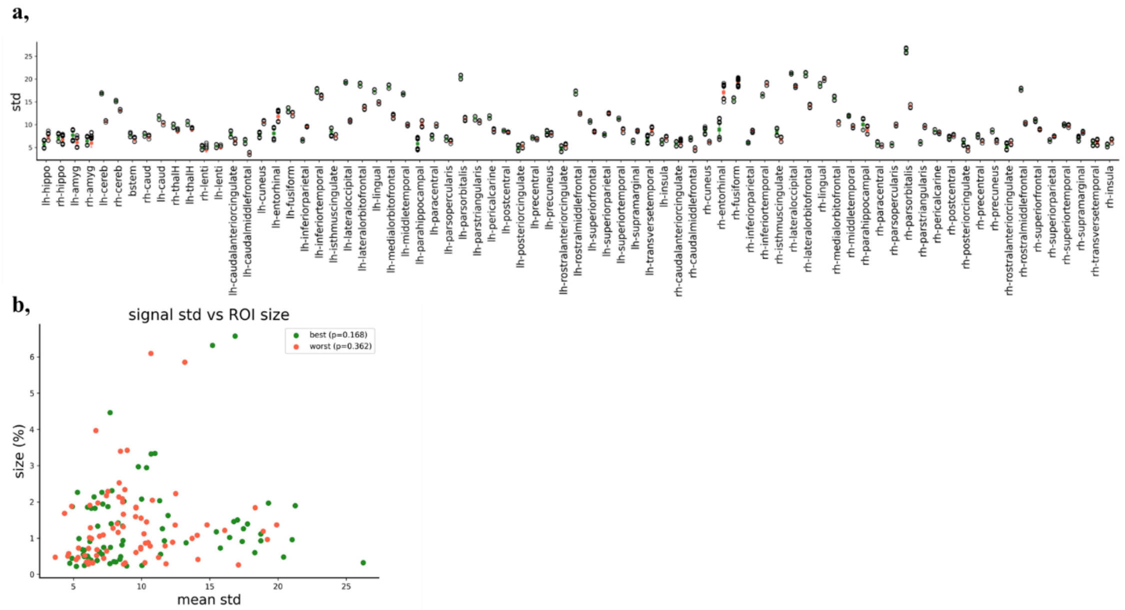


**Supp. Figure 2.3. a,** Regional coherence of the rs-fMRI data for delineated ROIs assessed as the standard deviation of the functional signal within the given region. The ROI naming conventions and assignment to lobes is detailed in *Supp. Figure 2.2.* **b,** Scatter plot of ROI's signal standard deviation with its size (as percentage of the whole GM) showing no apparent relationship between ROI signal homogeneity with size in our study. This suggests that in the analyses aiming to 'equalize' the regions within functional analyses, it might be preferable to sample the regions based on their signal variability rather than size.

#### *SI3. Univariate analyses of GM microstructure*

The univariate differences between infant groups in terms of the global microstructural features (i.e., median diffusion metrics across all cortical and subcortical ROIs detailed in *Supp. Figure 2.2.*) were assessed with paired t-tests corrected for multiple comparisons across groups and metrics using the Benjamini–Hochberg false discovery rate (FDR) correction. As expected, the three infant groups (PT:ses1, PT:ses2 and FT) differed significantly in terms of their global median diffusion characteristics, except for ODI in PT:ses2 vs FT comparisons (results not shown). Additionally, the global median metrics showed expected evolution with PMA at scan with visible different trends between groups (*Supp. Figure 3.1.*). All DTI-derived metrics (AD, RD, MD and FA) decreased with PMA at scan, with FA (and of lesser extent RD) showing the biphasic relationship with age previously described in Batalle et al. (2019). Both NODDI-derived metrics (ODI and NDI) increased with age suggesting the increasing complexity of the underlying microstructure.

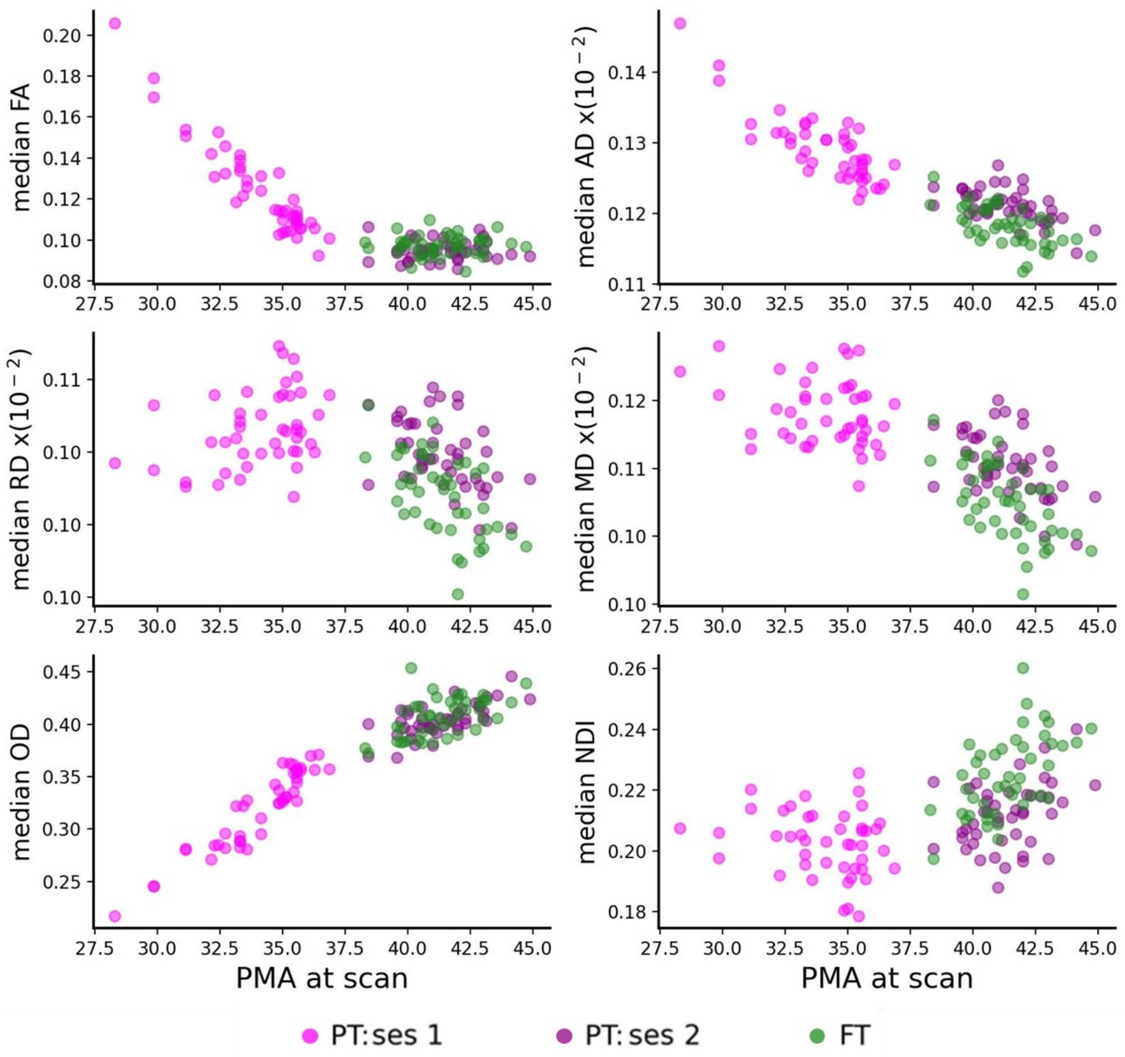


**Supp. Figure 3.1.** Relationship of median GM diffusion metrics with PMA at scan for the three infant groups: preterms at session 1 (PT:ses1, magenta), preterms at session 2 (PT:ses2, dark magenta), and paired controls (FT, green).

Additionally, we performed ANOVA modelling to investigate the effect of infant group, ROI and global median diffusion metrics, together with their interactions, on the regional microstructural features. The analysis revealed significant effects of these variables on the evaluated diffusion metrics (*Supp. Table 3.1.),* suggesting the existence of inter-regional differences in terms of their microstructural properties across subjects. ROI-specific diffusion metrics across infant groups are presented in the *Supp. Figure 3.2.*

**Supp. Table 3.1. a,** ANOVA model studying effects of PMA at scan and GA at birth on the six global median diffusion metrics. **b,** ANOVA model studying effect of ROI, infant group and residual global median diffusion and their interactions on each diffusion metric. p-value ≤0.001 [***], ≤0.01 [**], < 0.05 [*], <0.1 [.], ≥0.1 [*n.s.*].

**a,**

| | **Global FA** | **Global AD** | **Global RD** | **Global MD** | **Global OD** | **Global NDI** |
|---|---|---|---|---|---|---|
| *PMA scan* | *** | *** | *** | *** | *** | *** |
| *GA birth* | *** | *** | *** | *** | *** | *** |
| *PMA at scan: GA at birth* | *** | *** | *** | *** | *** | *** |

**b,**

| | FA | AD | RD | MD | OD | NDI |
|---|---|---|---|---|---|---|
| *ROI* | *** | *** | *** | *** | *** | *** |
| *Group* | *** | *** | *** | *** | *** | *** |
| *Global GM¤* | *** | *** | *** | *** | *** | *** |
| *ROI: Group* | *** | *** | *** | *** | *** | *** |
| *ROI: Global GM¤* | *n.s.* | *n.s.* | *** | *** | *** | *** |
| *Group: Global GM¤* | *n.s.* | *n.s.* | * | *n.s.* | *n.s.* | *n.s.* |
| *ROI: Group: Global GM¤* | *n.s.* | *n.s.* | *n.s.* | *n.s.* | *n.s.* | *n.s.* |

¤residual global median diffusion metric after correction for PMA at scan and GA at birth.

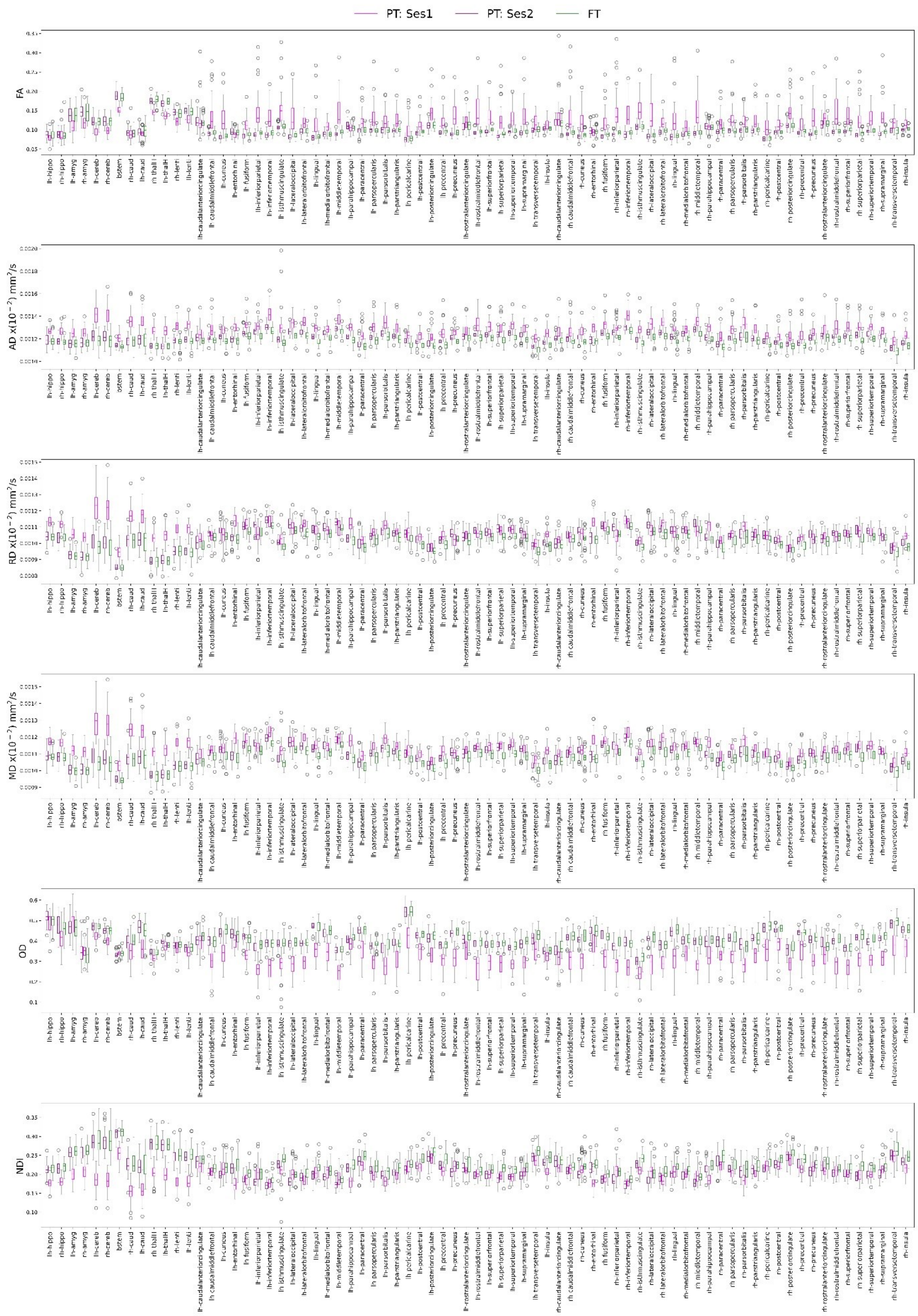


**Supp. Figure 3.2.** Diffusion metrics across ROIs and infant groups: preterm at session 1 (PT:ses1, magenta), preterm at session 2 (PT:ses2, dark magenta), and paired controls (FT, green). The ROI naming conventions and assignment to lobes is detailed in *Supp. Figure 2.2*.

### SI4. *Microstructural connectivity (MC) in infant groups: whole-brain analysis*

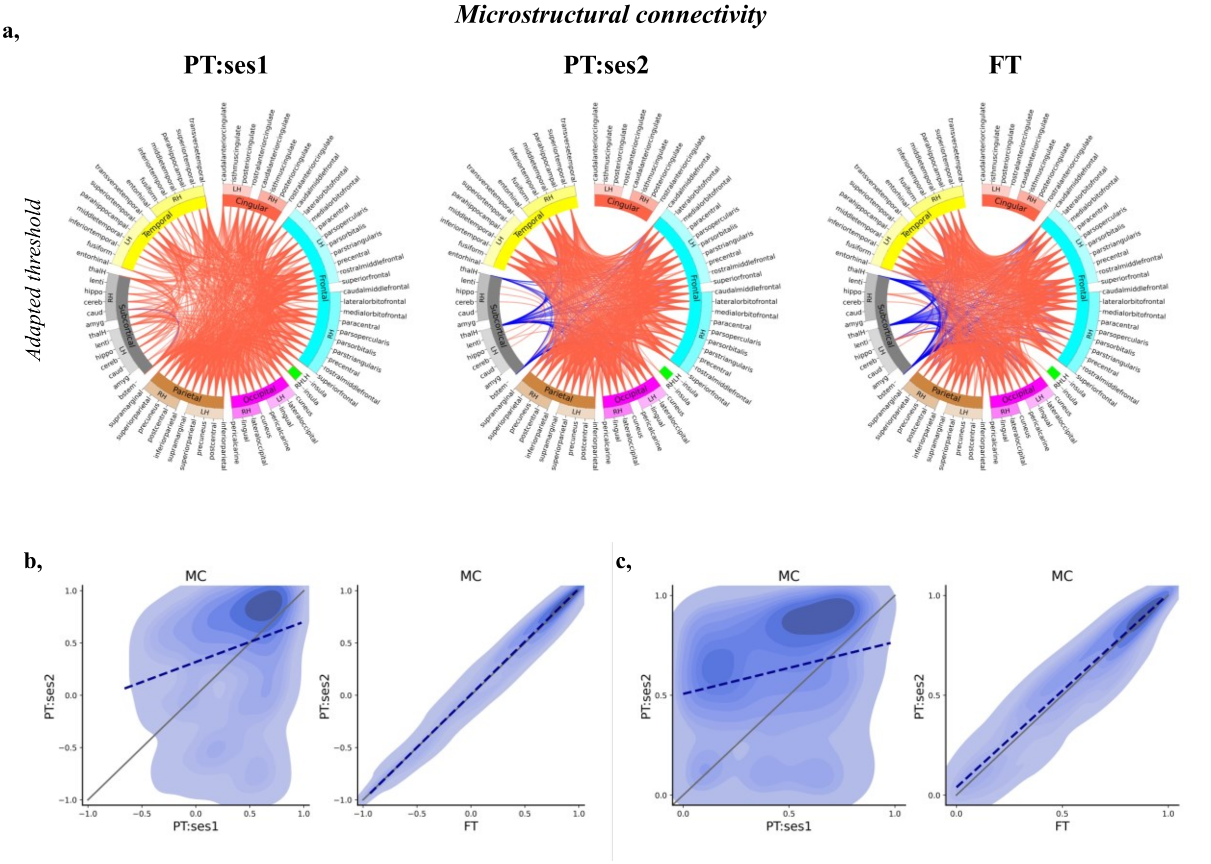


**Supp. Figure 4.1.** Microstructural connectome in infant groups. **a,** Circos plots representing correlation matrices and visualizing top 25 % MC connections for each infant group using adapted threshold of 0.657 for PT:ses1, 0.856 for PT:ses2, and 0.833 for FT. For the ease of visualization, cortical ROIs were grouped into 6 lobes (frontal in light blue, parietal in brown, temporal in yellow, occipital in pink, cingular in red, and insular in green) and subcortical ROIs (in grey) (see *Supp. Figure 2.2.* for ROI naming conventions and assignment to lobes). Connections with positive correlations are shown in red, negative in blue. **b,** Density plots showing relationships between the raw MC ROI pair similarity between PT:ses1 vs PT:ses2 (**left**) and FT vs PT:ses2 (**right**). The dotted blue line shows the significant* relationship determined by robust linear regression, while the solid grey line represents the identity relationship. **c,** Same visualisation as in (**b**) presenting results for absolute MC values. *after Benjamini–Hochberg false-discovery rate correction.

### SI5. *Evaluating functional connectivity (FC) in infant groups*

In the following section, we present results of the whole-brain evaluation of the FC for the 3 infant groups as well as analysis of the different connection subsets.

Similarly to MC, FC matrices were defined for each infant group, as the average of the temporal correlations of median functional signal between pairs of ROIs (*Figure 1.*) (note that the median was used to decrease the bias of the FC estimates for regions with high signal variability across voxels (*Supp. Figure 2.3.*)). Visualizing the strongest 25% connections, we observed a global trend towards reinforcement of the correlations across ROI connections with development (PT:ses1 vs PT:ses2 and FT) (*Supp. Figure 5a,b*). This was prominent for cortico-subcortical connections, maybe due to the higher cortico-cortical FC in the preterm period compared to later ages when synchronization of subcortical and cortical activity becomes more important, but it might also relate to methodological considerations (see *Discussion*).

***Functional connectivity***

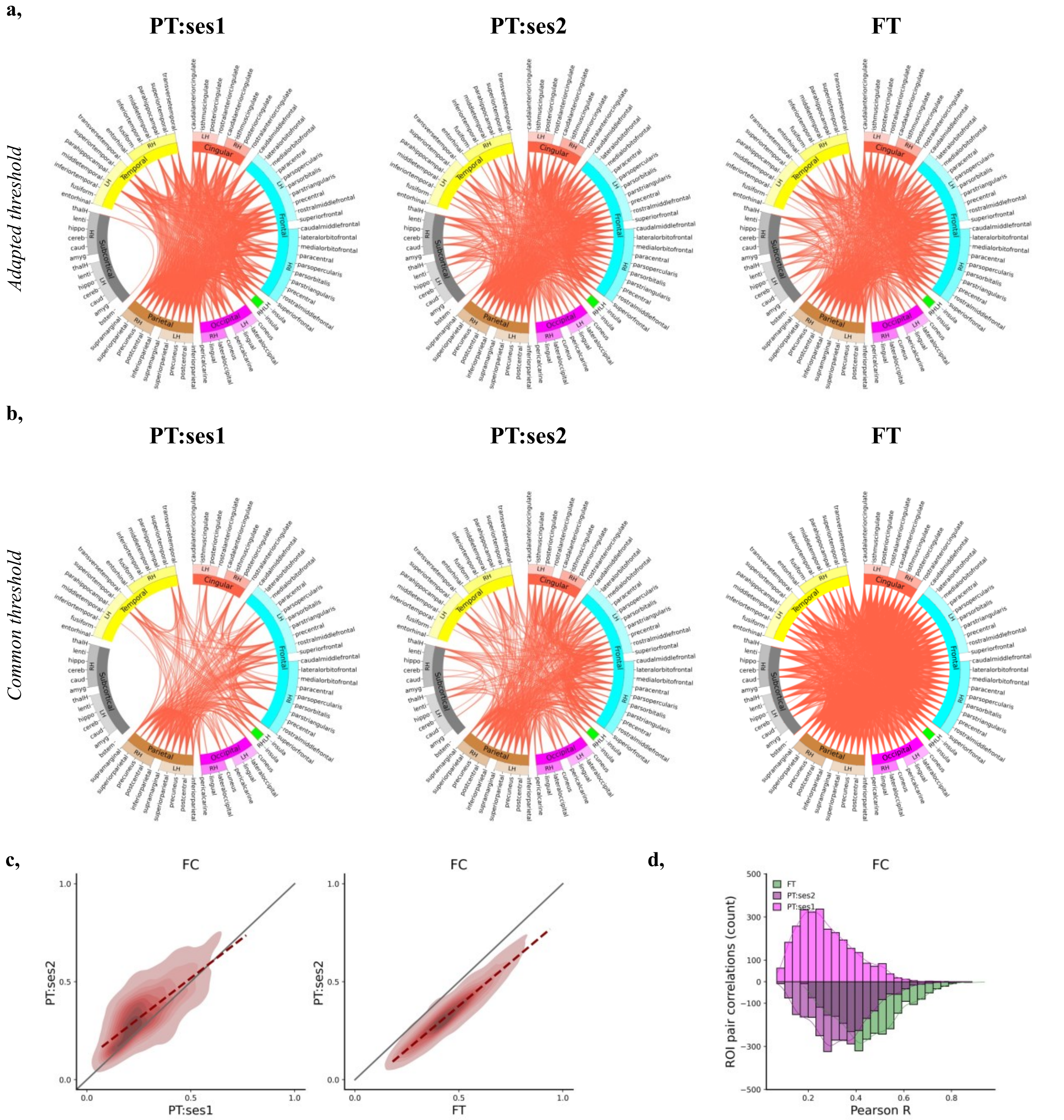

**e,**

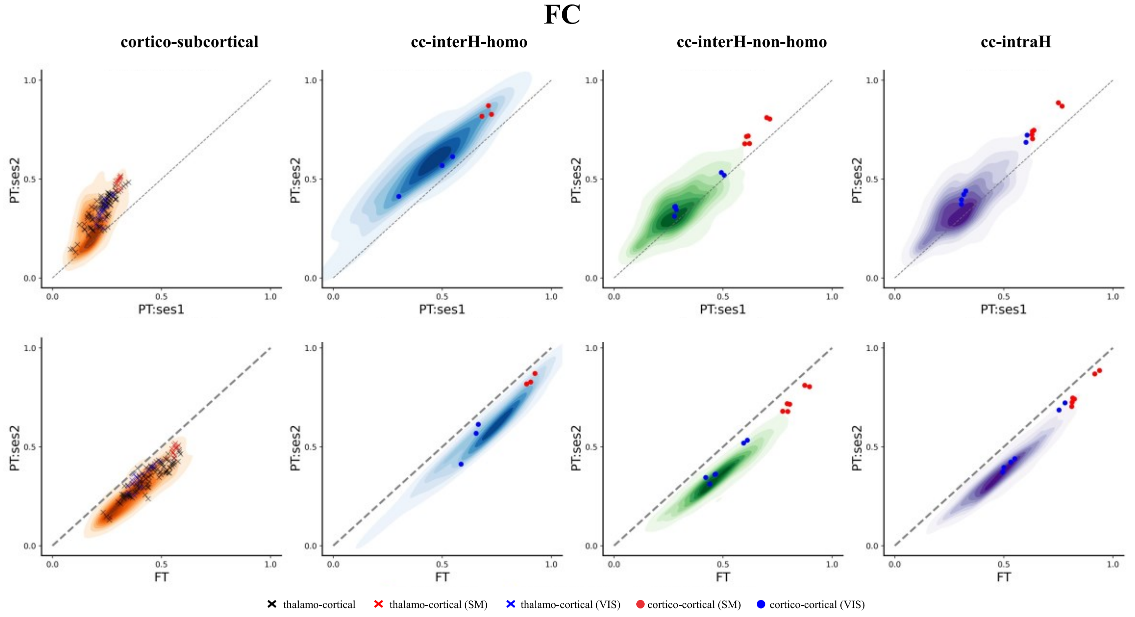


**Supp. Figure 5.1.** Functional connectome in infant groups. **a,** Circos plots representing correlation matrices and visualizing top 25 % FC connections for each infant group using adapted threshold of 0.349 for PT:ses1, 0.409 for PT:ses2, and 0.537 for FT. For the ease of visualization, cortical ROIs were grouped into 6 lobes (frontal in light blue, parietal in brown, temporal in yellow, occipital in pink, cingular in red, and insular in green) and subcortical ROIs (in grey) (see *Supp. Figure 2.2.* for ROI naming conventions and assignment to lobes). Connections with positive correlations are shown in red, negative in blue. **b,** Circos plots visualizing top 25 % FC connections for each infant group using a common FC threshold of 0.448. **c,** Density plots relationships between the FC ROI pair correlations between PT:ses1 vs PT:ses2 (**left**) and FT vs PT:ses2 (**right**). The dotted red line shows the significant relationship determined by robust linear regression, while the solid grey line represents the identity relationship. **d,** Distribution of ROI connection correlations (Pearson coefficient R) across infant groups. **e,** Changes of FC for subsets of ROI pairs between PT:ses1 vs PT:ses2 (top) and FT vs PT:ses2 (bottom). See *Figure 2* and *Supp. Figure 2.2.* for legend and colour codes.

As for MC, significant differences were observed in terms of correlation strengths across ROI connections with age suggesting ongoing development of FC during the preterm period (paired Wilcoxon test corrected for multiple comparisons: PT:ses2 > PT:ses1, W= 562652, $p<0.001$). Additionally, we observed a significant lowering of FC at TEA related to prematurity (PT:ses2 < FT, W= 125, $p<0.001$) (*Supp. Figure 5c*). Correlation strength was again highly related between PT:ses2 and FT across ROI connections (*Supp. Figure 5d*). In general, FC similarity between PT:ses1 and PT:ses2 groups was higher in comparison to MC (MC vs FC: Z=-22.22, $p<0.001$). This could suggest that early immature patterns of FC evolve during this period in a global and non-specific manner, while more specific changes in FC then take place at later stages.

To decipher potential FC differences related to maturational asynchrony, we again considered specific subsets of connections (*Supp. Figure 5e*). Cortico-subcortical FC showed globally lower strengths at PT:ses1, although it was increased at PT:ses2. Across cortico-cortical pairs, inter-hemispheric homotopic connections showed variable but globally higher FC strengths than cortico-subcortical connections at PT:ses1 and a rather homogenous increase in PT:ses2. Other cortico-cortical connections showed relatively low FC connectivity at PT:ses1 and limited change in PT:ses2. As with MC, FC in the primary SM system (and to a lesser extent the VIS system) showed higher values in all subsets of connections. A systematic reduction in FC strengths between PT:ses2 and FT was observed for all types of connections, suggesting a general decrease in FC linked to prematurity.

#### *SI6. Group-wise MC-FC analyses*

**a,**

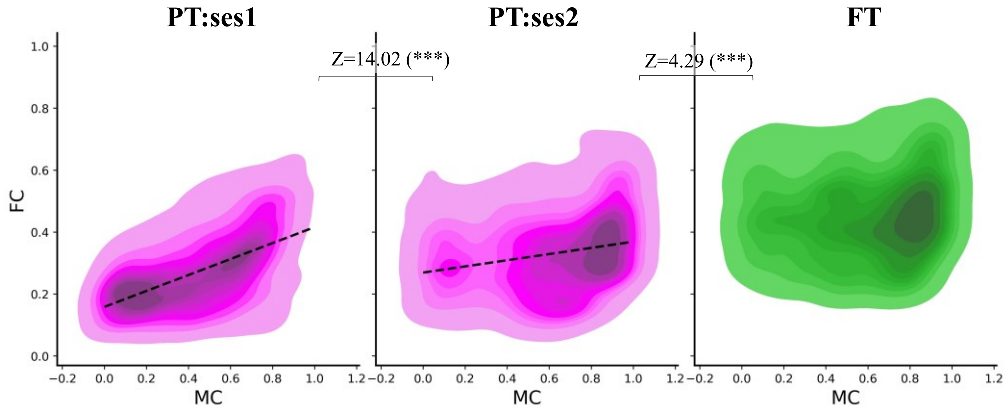


| | R | robust LR | permuted p |
|---|---|---|---|
| **PT:ses1** | 0.55 | 0.26x + 0.16 | 0.001 |
| **PT:ses2** | 0.24 | 0.10x + 0.27 | 0.013 |
| **FT** | 0.10 | 0.05x + 0.42 | 0.134 |

**b,**

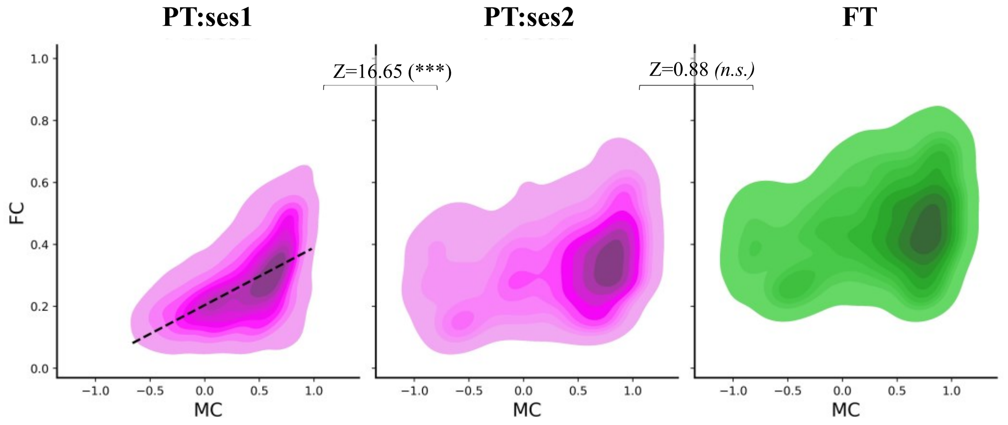


| | R | robust LR | permuted p |
|---|---|---|---|
| **PT:ses1** | 0.57 | 0.19x + 0.20 | 0.001 |
| **PT:ses2** | 0.32 | 0.07x + 0.30 | 0.188 |
| **FT** | 0.27 | 0.07x + 0.42 | 0.170 |

c,

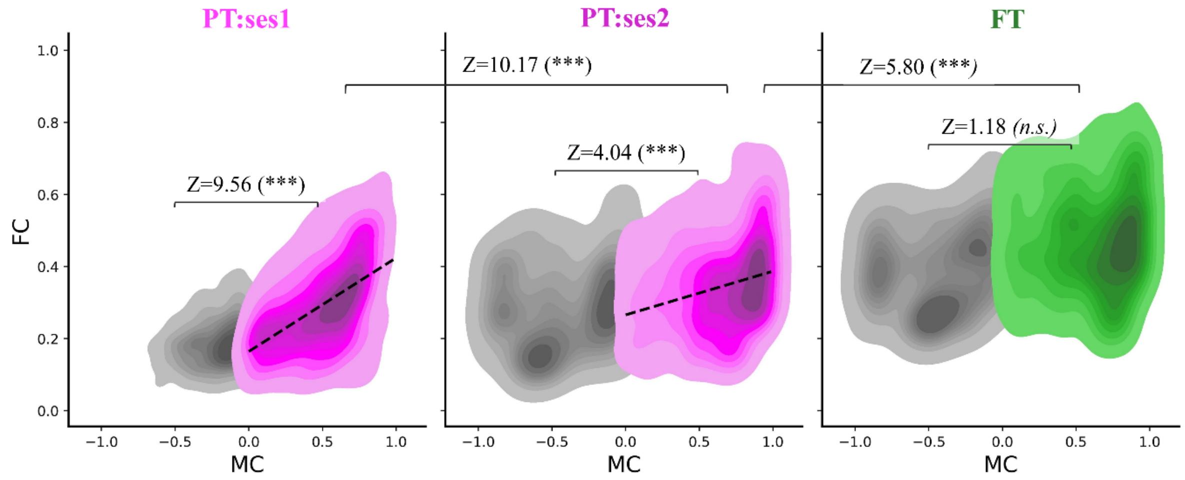


| | | R | robust LR | permuted p |
|---|---|---|---|---|
| **PT:ses1** | *positive* | 0.53 | 0.26x + 0.16 | 0.001 |
| | *negative* | 0.21 | 0.07x + 0.20 | 0.269 |
| **PT:ses2** | *positive* | 0.26 | 0.12x + 0.27 | 0.003 |
| | *negative* | 0.13 | 0.05x + 0.30 | 0.312 |
| **FT** | *positive* | 0.07 | 0.04x + 0.44 | 0.207 |
| | *negative* | 0.16 | 0.06x + 0.41 | 0.267 |

d,

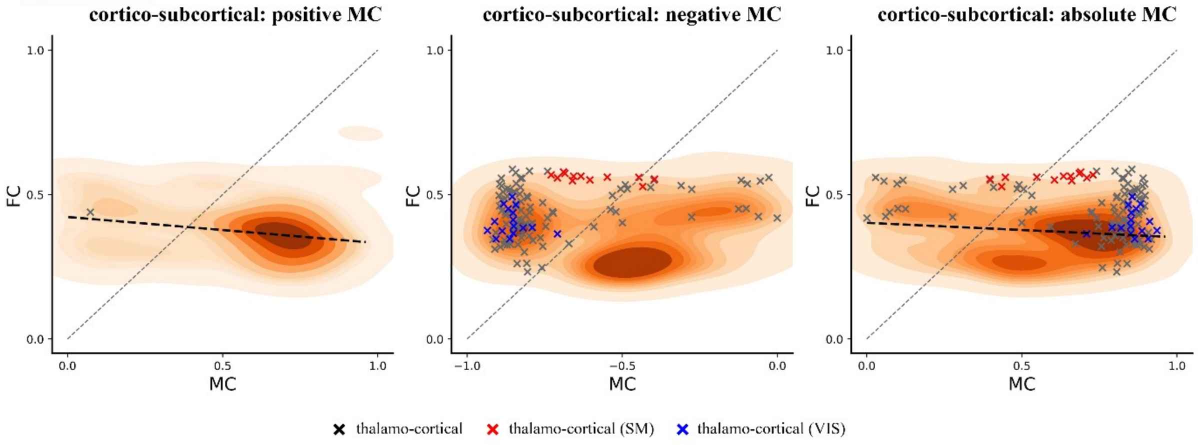


**Supp. Figure 6.1.** Relationship between MC and FC connectivity per ROI pair (upper triangle of symmetric matrices). **a,** MC-FC relationship after considering absolute values for the MC relationships. Same evaluation considering raw MC values is shown in (**b**) and considering ROI pairs with positive and negative MC values separately is shown in (**c**). The black dotted lines show the significant relationship determined by robust linear regression after permutation analysis. Tables summarize features of the robust linear relationship (LR) between the two connectomes. p-values $< 0.001$ ***, $p < 0.01$ **, $p < 0.05$ *, $p \geq 0.05$ *n.s.*. Z: Z-score comparing the slopes of linear regression. **d,** Relationship between MC and FC cortico-subcortical connections. Connections with positive (left) and negative (middle) MC in FT group, and relationship after taking their absolute values are shown for comparison. Significant negative relationships were found only in connections with positive connectivity strengths, suggesting different connectivity characteristics for cortico-subcortical vs cortico-cortical connectivity. The dotted black line shows the significant relationships determined by robust linear regression, while the grey line represents the identity relationship.

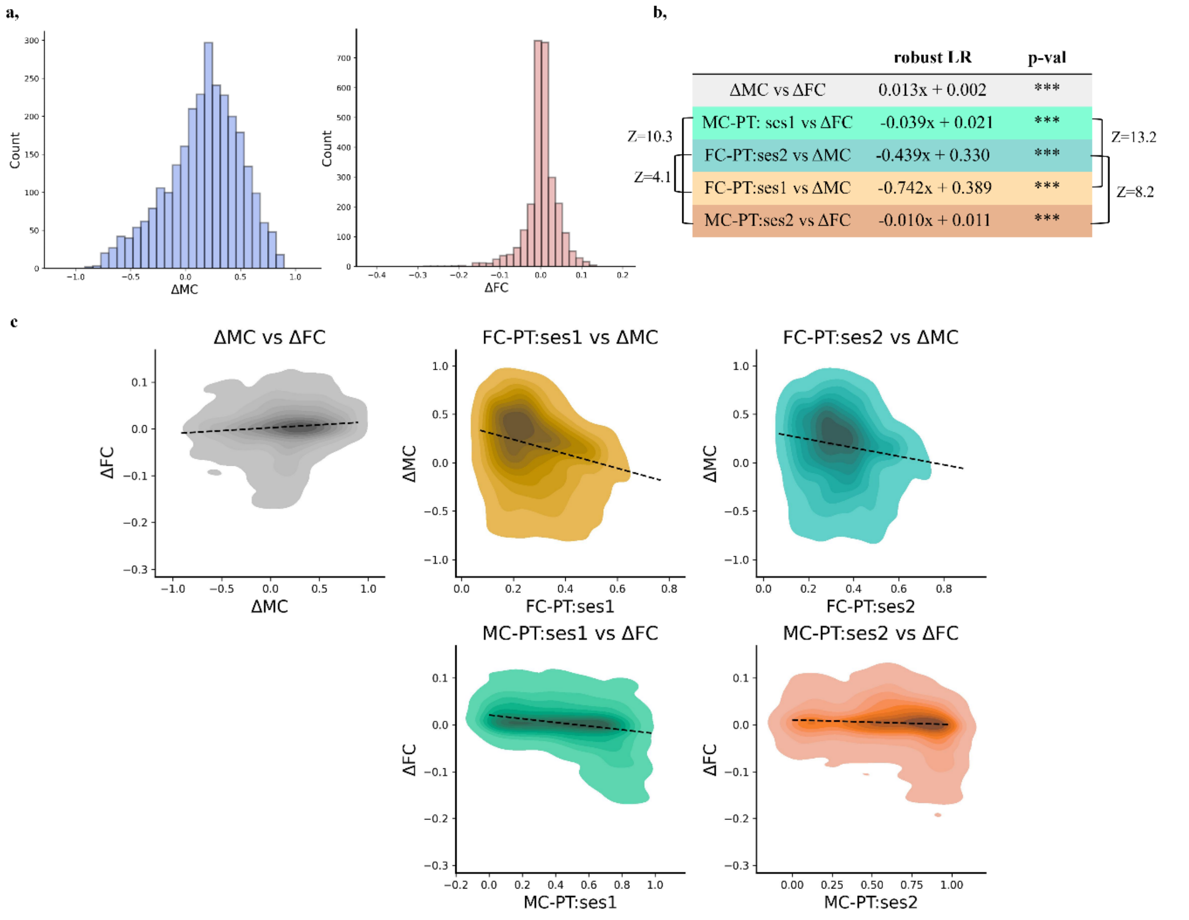


**Supp. Figure 6.2. a,** Distribution of changes in MC and FC strength between ROI pairs with age in preterm infants. **b,** The table summarizes relationships between the two connectomes estimated with robust linear regression (LR) (p-value *** <0.001 corrected for multiple comparisons). Z: Z-score comparing the estimated slopes. **c,** Relationship between ΔMC and ΔFC connectivity per ROI pair as well as the comparison to the connectivities of the opposite modality at different sessions. The black dotted lines show the significant relationship determined by robust linear regression. We colour coded table (**b**) and the subplots (**c**) by hypothesis tested by the given comparison: MC and FC co-evolve (grey), MC drives FC (cyan, dark cyan), and FC drives MC (orange, dark orange) presented in *Figure 5* in the main body of the manuscript.

The direct comparison between ΔMC and ΔFC suggested a low but significant positive relationship across ROI connections (robust linear regression: slope=0.01, p<0.001), suggesting some co-evolution of microstructural and functional connectomes. Besides, the developmental change in each modality was significantly related to the connectivity in the other modality at early session (ΔMC vs FC-PT:ses1: slope -0.74, p<0.001; ΔFC vs MC-PT:ses1: slope -0.04, p<0.001), but the slope of the relationship was significantly higher for ΔMC than ΔFC (Z=13.2, p<0.001), hinting at a potential higher dependency of FC development on the early underlying MC than the reverse. Comparing the ΔMC and ΔFC to the other modality at a later session suggested similar trend, with significant microstructural-functional relationships in both directions (MC vs FC-PT:ses2: slope -0.44, p<0.001; ΔFC vs MC-PT:ses2: slope -0.01, p<0.001) but again with a higher slope for ΔMC than ΔFC (Z=8.2, p<0.001). Finally, the relationships seemed stronger when comparing the evolution of MC and FC to the other modality at session 1 than at session 2 (ΔMC - FC-PT:ses1 vs FC-PT:ses2: Z=4.1; ΔFC - MC-

PT:ses1 vs MC-PT:ses2: Z=10.3), suggesting that changes in both MC and FC rely more on the early connectivity organization than they impact the later one.

### SI7. Network analyses

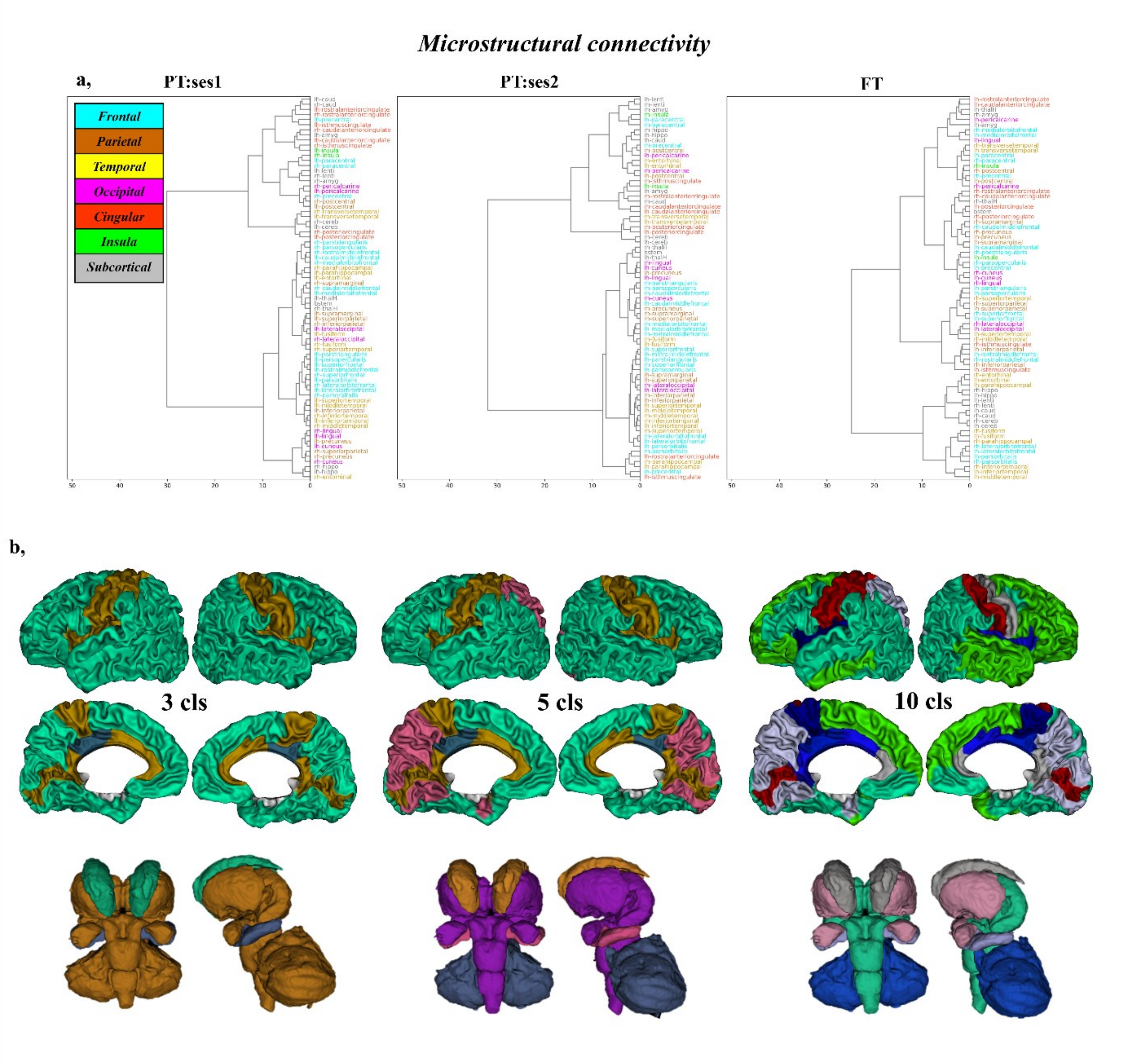


**Supp. Figure 7.1. a,** MC dendrograms for the three infant groups. The ROI naming conventions and assignment to lobes is detailed in *Supp. Figure 2.2.* **b,** Example representations clustering results in FT group for selected cluster umbers. The cluster colours are specific to this figure.

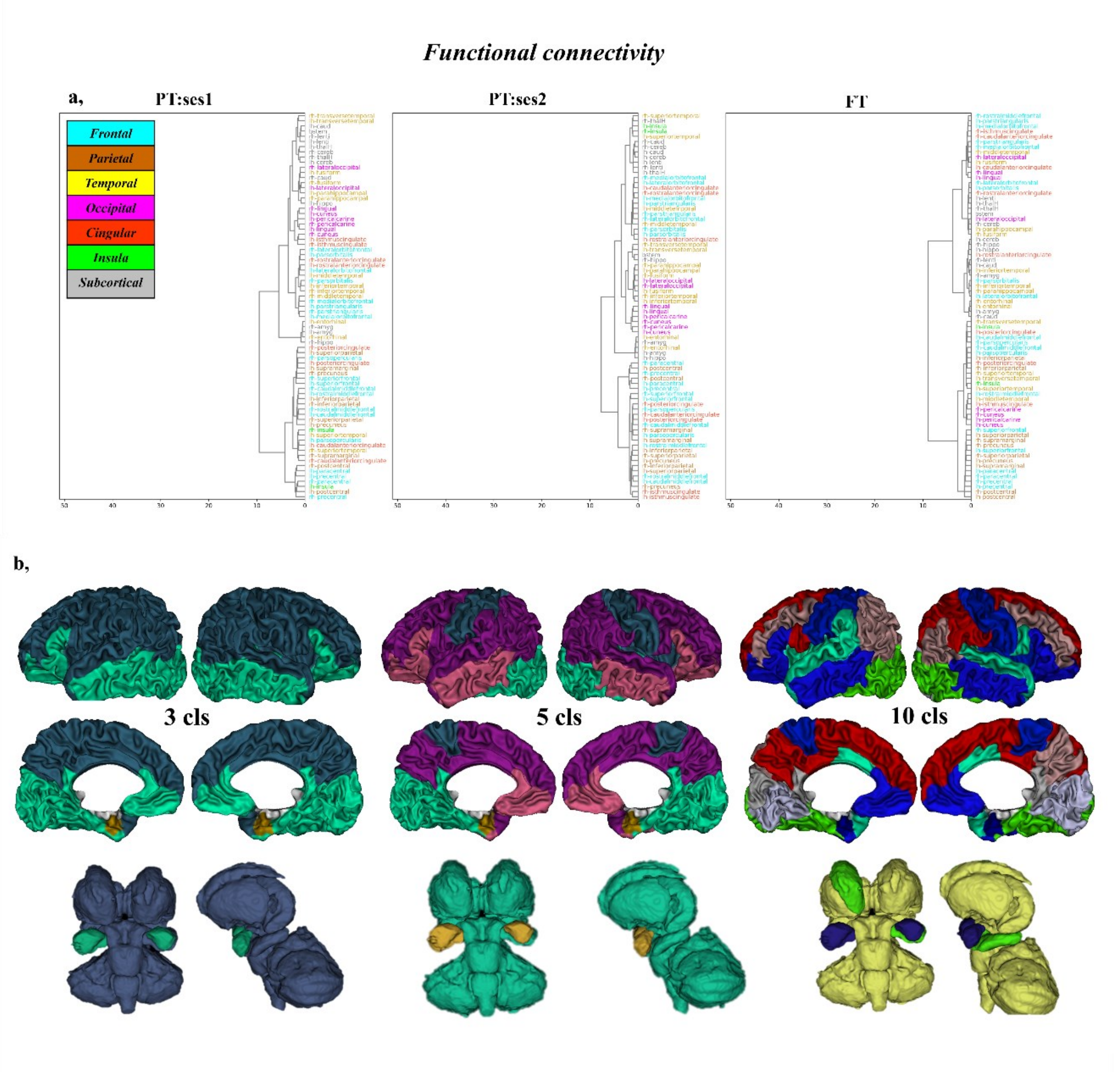


**Supp. Figure 7.2. a,** FC dendrograms for the three infant groups. The ROI naming conventions and assignment to lobes is detailed in *Supp. Figure 2.2.* **b,** Example of clustering results in FT group for selected cluster numbers. The cluster colours are specific to this figure.

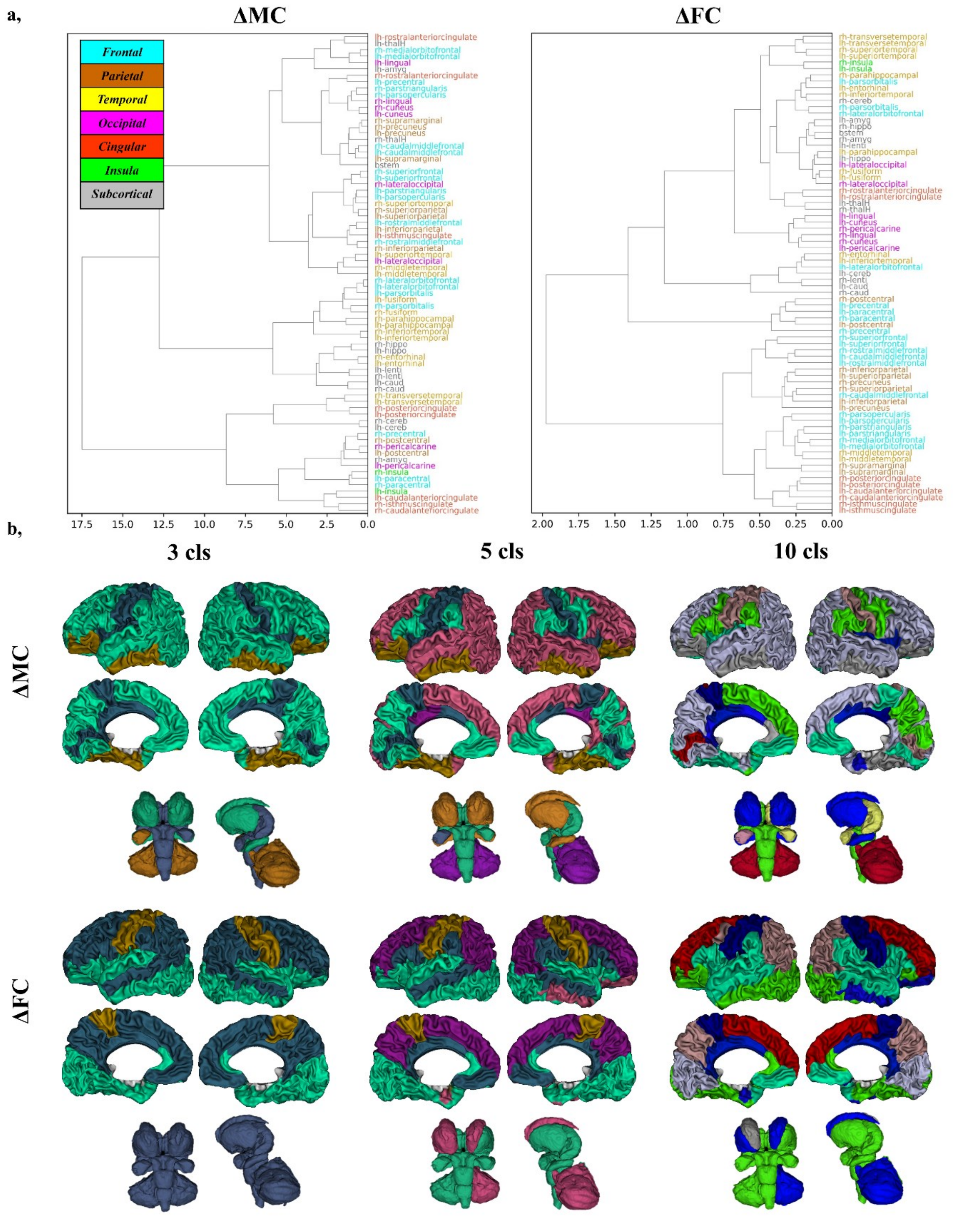


**Supp. Figure 7.3. a,** Dendrograms derived by clustering the ΔMC (**left**) and ΔFC matrices (**right**) developmental changes in preterm infants, between PT:ses1 and PT:ses2. The ROI naming conventions and assignment to lobes is detailed in *Supp. Figure 2.2.* **b,** Representations of example clustering results for selected cluster numbers for the two modalities. The cluster colours are specific to each figure.
